# Expression of TRPM_PZQ_ in excitable cells and sensory structures throughout the *Schistosoma mansoni* lifecycle

**DOI:** 10.64898/2026.10.16.759707

**Authors:** Claudia M. Rohr, Frances Blow, Sang-Kyu Park, Matthew Berriman, Jonathan S. Marchant

## Abstract

Praziquantel (PZQ) is an essential medicine used to treat schistosomiasis and other infectious diseases caused by parasitic flatworms. The parasite target of PZQ is a transient receptor potential ion channel of the melastatin subfamily, TRPM_PZQ_. As little is currently known about the native role of TRPM_PZQ_ in schistosomes, we developed and validated TRPM_PZQ_ specific antibodies to immunolocalize TRPM_PZQ_ across various *S. mansoni* life cycle stages (adult worms, juvenile worms, miracidia and cercariae). TRPM_PZQ_ was expressed in muscle, nervous tissues, as well as various sensory structures on the surface of the parasite across all these life cycle stages. In adult male worms, TRPM_PZQ_ positive neurons innervated the oral and ventral suckers as well as sensory papillae covering the head of the worm. TRPM_PZQ_ was also present in papillae throughout the body, and within tubercles. In a *S. mansoni* strain selected in the laboratory for resistance to PZQ, reduced expression and immunostaining of TRPM_PZQ_ was apparent. In immature worms, which can recover from PZQ exposure, TRPM_PZQ_ expression was predominantly found within the musculature compared with the neuronal localization evident in adult worms. A developmental transition of TRPM_PZQ_ from muscle to nervous tissues during schistosome maturation was also supported from single cell RNAseq data. Because muscle buffers Ca²⁺ transients more effectively than neurons, the transition of TRPM_PZQ_ away from muscle during schistosome maturation may increase the sensitivity of adult male worms to PZQ.

## Introduction

Schistosomiasis is a neglected tropical disease caused by infection with parasitic flatworms of the *Schistosoma* genus (*S. mansoni*, *S. haematobium* and *S. japonicum* as causative agents for human disease). Chronic schistosomiasis causes a significant health and financial burden, imposing considerable morbidity, mortality and economic strain on countries where transmission is endemic (1, 2).

The lifecycle of *Schistosoma* parasites requires passage through a vertebrate and an invertebrate (snail) host. Schistosome eggs excreted from an infected mammalian host hatch in fresh water to release free swimming miracidia. These motile larvae invade aquatic snails (the ‘intermediate’ host), where they undergo asexual reproduction. Once patent, infected snails release thousands of cercariae into the water, a second free swimming larval form. Cercariae penetrate through the skin of a mammal (the ‘definitive’ host), where they then develop into schistosomula. These larvae migrate through the dermis into the circulatory system reaching the lungs and then the liver where they develop into juvenile worms. After further maturation and migration into the mesenteric blood vessels (in the case of *S. mansoni* and *S. japonicum*), pairs of adult worms begin egg laying thereby completing the lifecycle as these eggs are excreted from host tissues.

Current treatment approaches depend on mass drug administration of praziquantel (PZQ), an old drug approved for use in the 1980s (3). PZQ exposure causes a rapid Ca^2+^ influx, spastic paralysis, and damage to the worm surface (tegument) which causes antigen exposure thereby triggering the elimination of parasites *in vivo* (3, 4). While PZQ is overall an excellent therapy offering cheap, safe and effective treatment, it also possesses some deficiencies. Most notably, PZQ is less effective at killing juvenile worms. While PZQ causes juvenile worms to rapidly contract, the immature worms exhibit a higher capacity for recovery. As such, there is a marked difference in the sensitivity of immature and adult worms to PZQ in measurements that encompass resilience (for example, a 30-fold difference in ED_50_ for worm reduction in a mouse model (5)).

This tolerance of juvenile worms toward PZQ presents a clinical challenge. Immature worm refractoriness to PZQ ensures that a single PZQ dose given to any patient may be non-curative, as juvenile worms that are not eliminated will mature into egg-laying adults thereby propagating the disease. Further, if someone is repeatedly infected, the probability of juvenile worms being present after treatment is even more likely (6, 7). Consequently, it is important to understand why juvenile worms display higher tolerance to PZQ in order to illuminate solutions for circumventing this challenge.

One potential mechanism for altering schistosome sensitivity to PZQ is through changes in the properties of the drug target. PZQ activates a non-selective cation channel in schistosomes, named TRPM_PZQ_ (8, 9), a member of the transient receptor potential melastatin subfamily. TRPM_PZQ_ is activated by (*R*)-PZQ in the sub-micromolar range, mimicking both the concentration and stereoselectivity of PZQ enantiomers active against adult *S. mansoni* (8, 9). While recent work has focused on characterizing the properties of TRPM_PZQ_ *in vitro* and validating this ion channel as the relevant clinical target of PZQ (reviewed in (10)), less is known about the endogenous properties of TRPM_PZQ_. Here, we developed custom antibodies to immunolocalize TRPM_PZQ_ across multiple life cycle stages of *S. mansoni*. Comparison between larval, immature and mature worms, as well as with a schistosome strain selected for PZQ resistance, highlighted changes in TRPM_PZQ_ expression and/or tissue localization that correlate with changes in the sensitivity of schistosomes to PZQ.

## Methods

### Antibody generation

Custom TRPM_PZQ_ (Smp_246790) antibodies were generated by Genscript (Piscataway, NJ). In brief, His-tagged constructs comprising either a NH_2_-(residues 1-402) or COOH-terminal region (residues 1709-1997) of *Sm*.TRPM_PZQ_ (8) were subcloned into the pET-30a vector and optimized for expression in *E. coli* (BL21 Star™, DE3). The COOH-terminal region overlaps with a similar region previously used to raise an antibody against *Schistosoma japonicum* TRPM_PZQ_ (*Sj*.TRPM_PZQ_, (11)).

Proteins were purified (from a 1L volume) using a nickel affinity column prior to immunization of two New Zealand rabbits per construct. The immunization protocol comprised three immunizations each two weeks apart, followed by small-scale antibody purification from individual rabbits to evaluate specificity, prior to an additional boost and then final antibody purification. A similar approach was used to generate a custom antibody targeting another TRP channel in the same subfamily, *Sm*.TRPM_MCLZ_ (Smp_333650,residues 1163-1380, (12)).

### Co-immunoprecipitation and mass spectrometry

The co-immunoprecipitation (co-IP) protocol was modified from a previously published method in schistosome worms (13). Briefly, adult worms (∼100 males) were homogenized using a Dounce homogenizer in 400 µl of ice-cold lysis buffer (50 mM NaCl, 5 mM EDTA, 1% Triton X-100, 20 mM Tris-HCl, pH 7.4,) supplemented with a protease inhibitor cocktail (Roche). The resulting homogenate was incubated at 4°C for 30 minutes with constant rotation to ensure complete lysis, followed by centrifugation (10,000 × g, 10 minutes at 4°C). The uppermost lipid layer was carefully removed, and the remaining clear supernatant was collected. A small aliquot (∼10 µl) was collected from this supernatant fraction and saved as an ‘input’ sample for Western blotting. The remaining supernatant was then incubated with a primary antibody (1 hour at 4°C) and then 20 µl of pre-washed protein A/G agarose beads (Roche) were added. Beads were previously prepared by washing with bead dilution buffer (130 mM NaCl, 0.05% Triton X-100, 0.1% lipid-free BSA, protease inhibitor cocktail, 10 mM Tris-HCl, pH 8.0) prior to resuspension in 200 µl of the same buffer. The bead-lysate mixture was incubated overnight (4°C with gentle rotation). Following incubation, the beads were washed three times with bead dilution buffer and then twice with Tris-saline buffer (140 mM NaCl, 10 mM Tris-HCl, pH 8.0).

The captured proteins were eluted by incubating the beads at 65°C for 5 minutes in an SDS-sample loading buffer. Control (IgG co-IP) and experimental (TRPM_PZQ_ co-IP) samples were processed by SDS-PAGE (4-12% Bis-Tris NuPage Mini-gel (Invitrogen) with a MOPS buffer system), and equivalent bands (∼250kDa) excised from the gel. Each band was processed by washing (25mM ammonium bicarbonate followed by acetonitrile), reduction (10mM dithiothreitol at 60°C followed by alkylation with 50mM iodoacetamide at room temperature), and then digestion with sequencing grade trypsin (Promega) at 37°C for 4h. The reactions were quenched with formic acid and the samples lyophilized.

Samples were then analyzed by nano LC-MS/MS (Waters M-Class LC system interfaced to a ThermoFisher Exploris 480 mass spectrometer) by MS Bioworks (Ann Arbor, Michigan). Peptides were loaded on a trapping column and eluted over a 75μm analytical column at 350nL/min. Both columns were packed with XSelect CSH C18 resin (Waters), the trapping column contained a 3.5μm particle, and the analytical column contained a 2.4μm particle. The column was heated to 55°C using a column heater (Sonation). The mass spectrometer was operated in data-dependent mode, with the Orbitrap operating at 60,000 FWHM and 15,000 FWHM for MS and MS/MS respectively. The instrument was run with a 3s cycle for MS and MS/MS. Data were searched using Mascot (Matrix Science). Mascot DAT files were parsed into Scaffold (Proteome Software) for validation, filtering and to create a non-redundant list per sample. Data were filtered at 1% protein and a peptide false discovery rate necessitating at least two unique peptides per protein.

### Western Blotting analysis

Proteins were resolved using SDS-PAGE (4–15% Tris-Glycine gradient gels, Bio-Rad) and transferred onto PVDF membranes using standard procedures. Membranes were blocked with 5% (w/v) bovine serum albumin (BSA, Millipore Sigma) in Tris-buffered saline supplemented with 0.1% Tween-20 (TBST). The membranes were then incubated overnight at 4°C with a custom TRPM_PZQ_ antibody (1:1000, Genscript) in 5% BSA-TBST. After washing with TBST, the membranes were incubated (1 h at room temperature) with a horseradish peroxidase (HRP)-conjugated secondary antibody (Invitrogen). Following additional washes, protein bands were visualized using a chemiluminescent substrate (ECL, Invitrogen) and imaged using a ChemiDoc™ MP Imaging System (Bio-Rad).

### S. mansoni life cycle stage isolation

All animal experiments followed ethical regulations approved by the Medical College of Wisconsin Institutional Animal Care an Use Committee. Juvenile and adult schistosomes were harvested from the liver and mesenteric vasculature of female Swiss Webster mice 16-49 days after infection with *S. mansoni* cercariae (NMRI or LE (PZQ-ER or PZQ-ES) strains (14)) performed at the Schistosomiasis Resource Center (Biomedical Research Institute, Rockville, MD).

Harvested schistosomes were washed in DMEM supplemented with penicillin-streptomycin (100 units/mL) and then transferred to ABC169 (15), for incubation overnight (37°C, 5% CO_2_) in vented petri dishes (100 x 25mm).

To obtain miracidia, mature eggs were resuspended in artificial pond water (0.46 µM FeCl_3_⋅6H_2_O, 220 µM CaCl 2H O_2_, 100 µM MgSO_2_⋅7 H O, phosphate buffer [313 µM KH_2_PO_4_, 14 µM (NH_4_)_2_SO_4_] pH 7.2) and exposed to light in a 35mm dish. *S. mansoni* cercariae were obtained from patent *Biomphalaria glabrata* snails (Schistosomiasis Resource Center) by exposing snails to direct light for 1 hour. Schistosomula were then generated via mechanical transformation of cercariae. To do this, a cercarial suspension in a 50mL conical tube was placed on ice and vortexed vigorously. The suspension was then layered onto a 70% Percoll gradient and centrifuged (15 minutes at 500 x g at 4°C). Percoll was pipetted out of the tube until cercaria heads became visible in this discard; at which point 5-10 mL of RPMI was added and centrifugation repeated (5 minutes at 100 x g, 4°C). This process was repeated until the majority of tails were removed and the heads remained in the gradient. Mechanically transformed cercaria were then pelleted, washed twice with RPMI and placed into a 6 well plate containing DMEM supplemented with 2% FBS & 2% Penicillin Streptomycin (Invitrogen), and incubated for 24 hours at 37°C, 5% CO_2_.

### Immunolabeling

Miracidia and cercariae were fixed as described previously (16). In brief, an 8% formaldehyde solution (37% formaldehyde diluted in artificial pond water) was added to an equal volume of artificial pond water containing either miracidia or cercariae. The solutions were mixed well and placed on a rocker at room temperature for 2 hours. Fixed samples were washed in PBSTx (1x PBS, supplemented with 0.3% Triton X-100) and then placed in PBS until staining. Schistosomula, juveniles, and adult worms were relaxed with 2.5% tricaine (ethyl 3-aminobenzoate methanesulfonate in PBS) for 2 minutes, then incubated in 0.6M MgCl_2_. Samples were fixed (4% formaldehyde in PBSTx) either overnight at 4°C, or for 4 hours on a rocker at room temperature. Following fixation, all samples were washed 3x in PBS. For subsequent steps, samples were transferred to a black walled clear bottom 96 well plate (for miracidia, cercaria, and schistosomula) or in the case of juveniles and adult worms to a 24 well plate containing 35 µm mesh baskets (Intavis).

All samples were permeabilized for 20 minutes (1% SDS + 1x PBS), rinsed in PBSTx and incubated in blocking solution (5% goat serum (Invitrogen), 0.5% Western Blocking Reagent (Millipore Sigma) in TNTx (0.1 M Tris pH 7.5, 0.15 M NaCl, and 0.1% Tween-20)) for 1-2 hours. Following blocking, samples were incubated overnight at 4°C in primary antibody (1:50-1:100) that was diluted in blocking solution. Primary antibodies were against TRPM_PZQ_ or TRPM_MCLZ_ (1:75), synapsin (DSHB, 3C11, 1:50) and β-tubulin (DSHB, E7, 1:100). The following day, samples were washed for ≥2 hours in PBSTx and incubated overnight in goat-anti-rabbit Alexa Fluor 568 (1:250) for TRPM_PZQ_ and

TRPM_MCLZ_, goat-anti mouse Alexa Fluor 488 (1:250) for synapsin and β-tubulin, DAPI (1 µg/mL in PBSTx), or Alexa Fluor 647 conjugated phalloidin (66nM final in PBSTx, Invitrogen) diluted in the blocking solution. Following this overnight incubation, samples were washed 2x in PBSTx and mounted on slides using Prolong Diamond Antifade Mountant (Invitrogen). For stimulated emission depletion (STED) microscopy, adults were fixed and incubated in primary antibody following the same methods detailed above. Following the >2 hour PBSTx wash, worms were incubated overnight in goat-anti-rabbit STAR ORANGE (1:250) (Abberior), goat-anti mouse Alexa Fluor 488 (1:250), DAPI and Alexa Fluor 647 conjugated phalloidin. Samples were washed 2x in PBSTx and mounted using Abberior Mount liquid antifade mountant.

### Image acquisition

Samples were imaged using either a Dragonfly 502 (Zyla 4.2 PLUS sCMOS camera) or Dragonfly 620SR (ZL41 Cell 4.2 sCMOS camera) (Oxford Instruments). Both spinning disc confocals were interfaced with inverted Leica DMi8 microscopes and all images were acquired using a 63x HC PL APO 1.4NA or 100x-TIRF HC PL APO 1.47 oil immersion objective (Leica). Images were captured and stitched using Fusion (Version 2.7, Oxford Instruments) and processed using Imaris (Version 11.0.0, Oxford Instruments). STED samples were imaged using the FACILITY line 3D-STED super-resolution imaging system (Abberior) housed on an inverted Evident Ix83 microscope. Images were acquired at 60x using the Lightbox Software (Version 2024.48.21878-gc86bbd647c, Abberior) and processed using Imaris.

### Image analysis

Image analysis was performed using ImageJ (version 1.54s). A maximum projection Z-projection of the full volume of the region of interest (e.g. individual tubercles or the surface of the sucker) was made to capture the entire structure. The image stack was normalized to background fluorescence captured in an image plane with no staining. For fluorescence profiles, a single ∼20 µm line was drawn spanning the region surrounding the structure of interest and the Plot Profile function was used to measure fluorescence intensity. For single tubercle imaging, ROIs were placed on the corpuscle or on the tubercle surface surrounding the corpuscle (20 tubercles from 2 different worms). Mean intensity was measured and the difference between the corpuscle and the tubercle background was calculated. Welch’s t-test was used to compare the means of fluorescence intensity resolved in different image stacks captured from PZQ-ER and PZQ-ES worms. Statistical analyses were performed using Prism (GraphPad, v. 11.0.0).

### Analysis of single cell RNA sequencing datasets

We used previously published single-cell RNASeq datasets to examine TRPM_PZQ_ expression in juvenile (17) and adult (18) muscle and neural cell types. Both datasets were analysed as outlined in the original publications. For the juvenile dataset the raw counts were downloaded and re-analysed following methods in the original publication. Cluster annotations were performed *de novo* as these were not included in the downloaded raw data. For the adult dataset, the analysed Seurat object from the original publication was downloaded, and published cell type annotations were used. For each dataset all cells assigned to muscle sub-types were combined into a “muscle” cell type and all cells assigned to neural sub-types were combined into a “neuron” cell type. We then examined TRPM_PZQ_ expression within these broader muscle and neuron cell types. “Percent expressed” refers to any cell with non-zero TRPM_PZQ_ expression after normalisation. To calculate scaled relative expression for each cell type, average expression values were min-max normalised to a range of 0 to 1 independently for each dataset, comparing all broad cell types in the dataset, such that the cell type with the lowest average expression was set to 0 and the highest to 1.

## Results

### Antibody Validation

Two different regions of *Schistosoma mansoni* TRPM_PZQ_ (*Sm*.TRPM_PZQ_) were targeted for polyclonal antibody generation. These comprised a NH_2_-terminal (1-402 amino acids, α-TRPM_PZQ_-N) or a COOH-terminal region (1709-1997 amino acids, α-TRPM_PZQ_-C) within *Sm*.TRPM_PZQ_ (**Figure 1A**). Both antibodies were profiled for target specificity using the following approaches.

**Figure 1.**
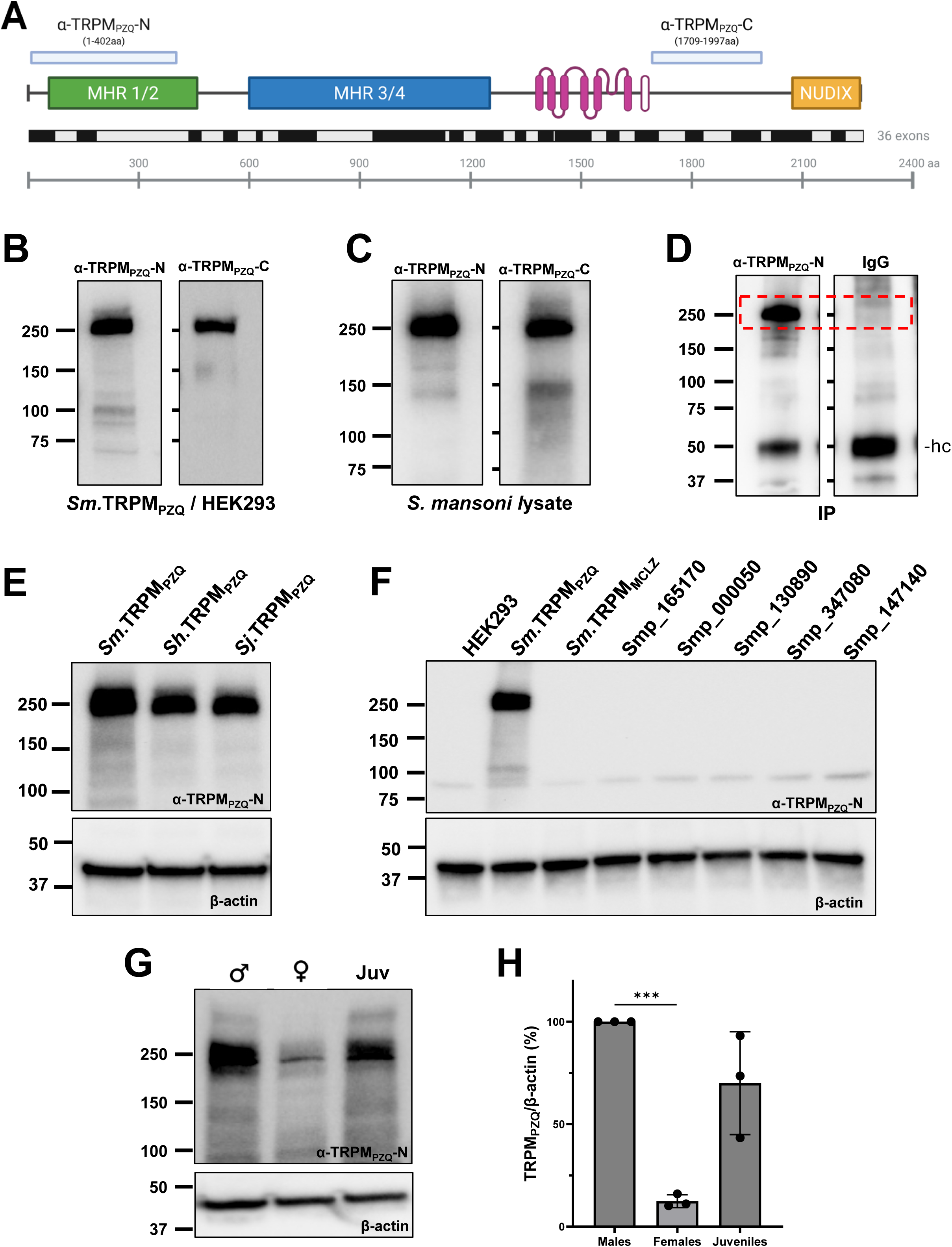
Validation of TRPM_PZQ_ antibodies. (**A**) Regions within TRPM_PZQ_ used to generate NH_2_-(α-TRPM_PZQ_-N) and COOH-(α-TRPM_PZQ_-C) antibodies. Schematic of TRPM_PZQ_ structures highlights: *green*: TRPM homology region 1/2; *blue*: TRPM homology region 3/4, *pink*: transmembrane domains (solid) and TRP helix (open), *yellow*: nudix hydrolase domain. Exon organization and an amino acid scale is shown. (**B**) Representative Western blot using NH_2_-(α-TRPM_PZQ_-N) and COOH-terminal (α-TRPM_PZQ_-C) TRPM_PZQ_ antibodies against a HEK293 cell line expressing *S. mansoni* TRPM_PZQ_ (*Sm*.TRPM_PZQ_). (**C**) Western blot using α-TRPM_PZQ_-N and α-TRPM_PZQ_-C antibodies against *S. mansoni* adult male worm lysate. (**D**) Western blot of native immunoprecipitation probed with α-TRPM_PZQ_-N and blotted for the COOH-terminal antibody. Dotted red box indicates the gel region excised and processed for mass spectrometry. ‘hc’ indicates heavy chain. (**E**) Western blot assessing cross reactivity of α-TRPM_PZQ_-N against TRPM_PZQ_ in other schistosome species: *S. japonicum* TRPM_PZQ_*, Sj*.TRPM_PZQ_; *S. haematobium* TRPM_PZQ_*, Sh*.TRPM_PZQ_. (**F**) Western blot using NH_2_-terminal TRPM_PZQ_ antibody of HEK293 cells expressing various annotated schistosome TRPM channels. (**G**) Representative Western blot of adult female, male and juvenile *S. mansoni* worm lysates using α-TRPM_PZQ_-N. (**H**) Densitometry from n=3 Western blots of the same life cycle stages shown in (G). Significance was determined for using a Welch’s t-test, *** p < 0.001.

First, in a stable HEK293 cell line induced to express *Sm.*TRPM_PZQ_, each antibody recognized a predominant band consistent with the predicted size of *Sm*.TRPM_PZQ_ (∼255kDa, **Figure 1B**).

Second, Western blots were performed using a crude lysate prepared from *S. mansoni* adult male worms. Again, both antibodies recognized a predominant ∼250kDa band (**Figure 1C**), consistent with the size of *Sm*.TRPM_PZQ_ observed after heterologous expression of *Sm*.TRPM_PZQ_ in HEK293 cells (Figure 1B). Some weaker staining of lower sized bands was also observed, more notably with the COOH terminal antibody (α-TRPM_PZQ_-C). Therefore, α-TRPM_PZQ_-N was used for subsequent experiments.

Third, we performed a native immunoprecipitation (IP) using the NH_2_-terminal antibody (α-TRPM_PZQ_-N) or a parallel IgG control. IP samples were separated by SDS-PAGE and bands around 255kDa were excised from both the *Sm.*TRPM_PZQ_ and IgG control lanes and analyzed by mass spectrometry (**Figure 1D**). After exclusion of proteins present in the IgG control band, the top proteins found in the TRPM_PZQ_ sample were spectrin β chain (*Smp_143470*), TRPM_PZQ_ (*Smp_246790*) and paramyosin (*Smp_0885540*). Peptide coverage for *Sm.*TRPM_PZQ_ is presented in Supplementary Figure 1.

Fourth, further Western blots were performed to screen for cross reactivity against other TRPM_PZQ_ orthologs and paralogs. The α-TRPM_PZQ_-N antibody recognized TRPM_PZQ_ in other schistosome species as both *S. haematobium* TRPM_PZQ_ (*Sh*.TRPM_PZQ_ (19)) and *S. japonicum* TRPM_PZQ_ (*Sj*.TRPM_PZQ_, (19)) were detected after heterologous expression (**Figure 1E**). However, when the same NH_2_-terminal antibody was screened against the other annotated schistosome TRPM channels that were heterologously expressed in HEK293 cells, no specific signal was resolved despite some of these TRPM channels displaying sequence homology in the NH_2_-terminal MHR region (**Figure 1F**).

Fifth, endogenous expression of TRPM_PZQ_ was examined in adult male, adult female and juvenile worms by Western blotting (**Figure 1G**). TRPM_PZQ_ expression was highest in adult males and lowest in adult female worms (**Figure 1H**), a result consistent with prior mRNA analysis and immunostaining (11, 20).

Finally, to better define the region of TRPM_PZQ_ recognized by α-TRPM_PZQ_-N, we performed exon by exon deletions of the NH_2_-terminal region of TRPM_PZQ_. Deletion of exon 4 caused a complete loss of α-TRPM_PZQ_-N immunoreactivity when these constructs were expressed in HEK293 cells (Supplementary Figure 2). This confirms recognition by α-TRPM_PZQ_-N of a region within the recombinant protein used for immunization.

Based on all these assessments which demonstrate specificity toward *Schistosoma spp.* TRPM_PZQ_, we used α-TRPM_PZQ_-N to examine the native localization of TRPM_PZQ_ across different *S. mansoni* life cycle stages.

### Adult worm

Immunofluorescence staining of *S. mansoni* adult male worms was imaged using confocal microscopy. TRPM_PZQ_ expression was evident in neuronal processes (co-stained with β-tubulin), muscle (co-stained with phalloidin) as well as various sensory surface structures present across the worm surface (**Figure 2A**). No staining was evident in the adult worm, or other life cycle stages, after incubation with the secondary antibody alone (Supplementary Figure 3).

**Figure 2.**
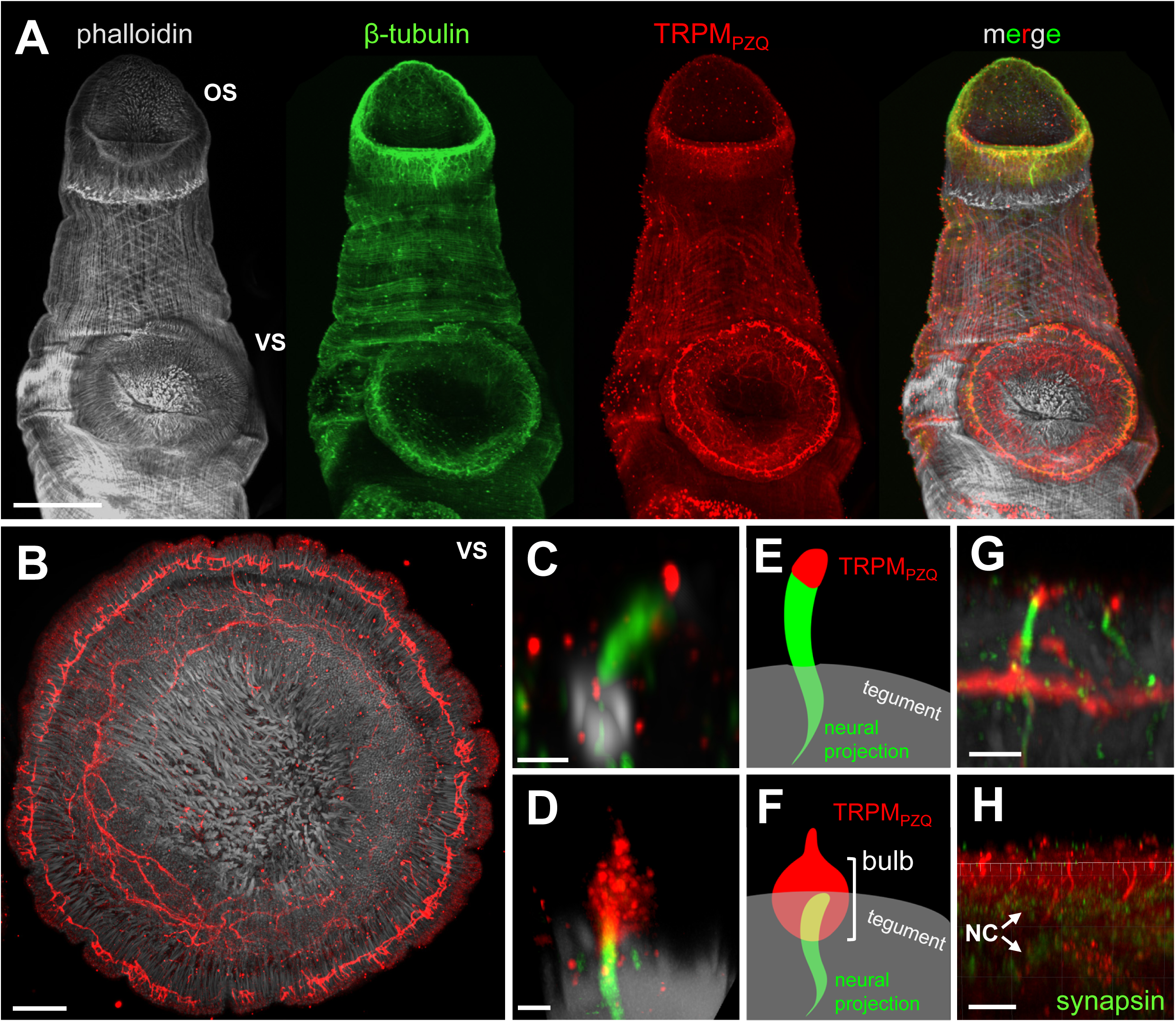
TRPM_PZQ_ staining within the head of adult *S. mansoni* worms. Unless otherwise noted, all samples show labeling of TRPM_PZQ_ (red), β-tubulin (green), phalloidin (grey). Abbreviations: oral sucker (‘OS’), ventral sucker (‘VS’), nerve cords (‘NC’). (**A**) Maximum intensity projections of staining of an adult (49 DPI) male *S. mansoni* worm using antibodies against phalloidin, β-tubulin and TRPM_PZQ_ (α-TRPM_PZQ_-N), scale bar: 100 μm. (**B**) Maximum intensity projection of the ventral sucker, stained for phalloidin and TRPM_PZQ_. Scale bar: 20 μm. (**C**) Digital-form papilla. 3D-STED imaging of a single digital-form papilla emerging from a pore on the worm surface. Scale bar: 1 µm. (**D**) Hemispherical papilla. 3D-STED imaging of hemispherical papilla. Scale bar: 1 µm. (**E-F**) Schematic depicting TRPM_PZQ_ localization in both types of papillae. (**G**) Axial (‘xz’) projection showing innervation of digital-form papillae from the neuronal ring surrounding the ventral sucker. Scale bar: 2 µm. (**H**) Axial (‘xz’) projection showing hemispherical papillae projecting down toward the nerve cords captured ventrally between the suckers. Scale bar: 10 μm.

TRPM_PZQ_ staining distribution in different excitable cell types is apparent from images of the head of the worm, encompassing both the suckers and the area between them (Figure 2A). Staining encompassed an elaborate ring of TRPM_PZQ_-positive neurons that runs alongside the circular muscle circumscribing the ventral sucker (**Figure 2B**, Supplementary Figure 4), staining of diagonal and longitudinal muscles in the region between the suckers (Figure 2A), and many punctate structures across the worm surface (Figure 2A). These punctate structures correspond to different types of sensory papillae that have been previously described in the literature (21, 22) and were found across the dorsal and ventral surfaces of the head. To obtain higher resolution images, STED microscopy was used to image individual structures evidencing two main types - “digital-form” papillae (a ‘finger-like’ projection, **Figure 2C**) and “hemispherical” papillae (comprising a surface ‘bulb’ and a cilium at the tip, **Figure 2D**). A schematic representation of these structures is depicted in **Figures 2E&F**. Both these types of papillae were spaced across the face of both suckers, but only hemispherical papillae were found in the region between the suckers. TRPM_PZQ_ localized to the distal end of the process within digital-form papillae and within both the ‘bulb’ and cilium within hemispherical papillae (Figure 2C-F). Nervous system innervation in both papillae types was evident supporting their sensory role(s). Neuronal processes projecting from the ventral sucker nerve ring interfaced with digital-form papillae (**Figure 2G**). Similarly, in the hemispherical papillae found between the suckers, TRPM_PZQ_ positive processes from the papillae projected deeper into the worm parenchyma (**Figure 2H**).

TRPM_PZQ_ expression was also evident in other sensory structures distributed across the body surface. This included staining of the aforementioned papillae but also included clear focal staining within tubercles found on the dorsal surface of male worms (**Figure 3A**). The tubercular staining pattern for TRPM_PZQ_ on the body of the worm was dramatic, with a punctate fluorescence spot evident at the center of each tubercle (Figure 3A). This ‘bullseye’ pattern resulted from staining of a structure known as the ‘sensory corpuscle’ which resides at a central localization within the tubercle (23). Validation of this staining was performed with both antibodies (α-TRPM_PZQ_-N and α-TRPM_PZQ_-C), which resulted in similar focal staining within the tubercle (Supplementary Figure 5).

**Figure 3.**
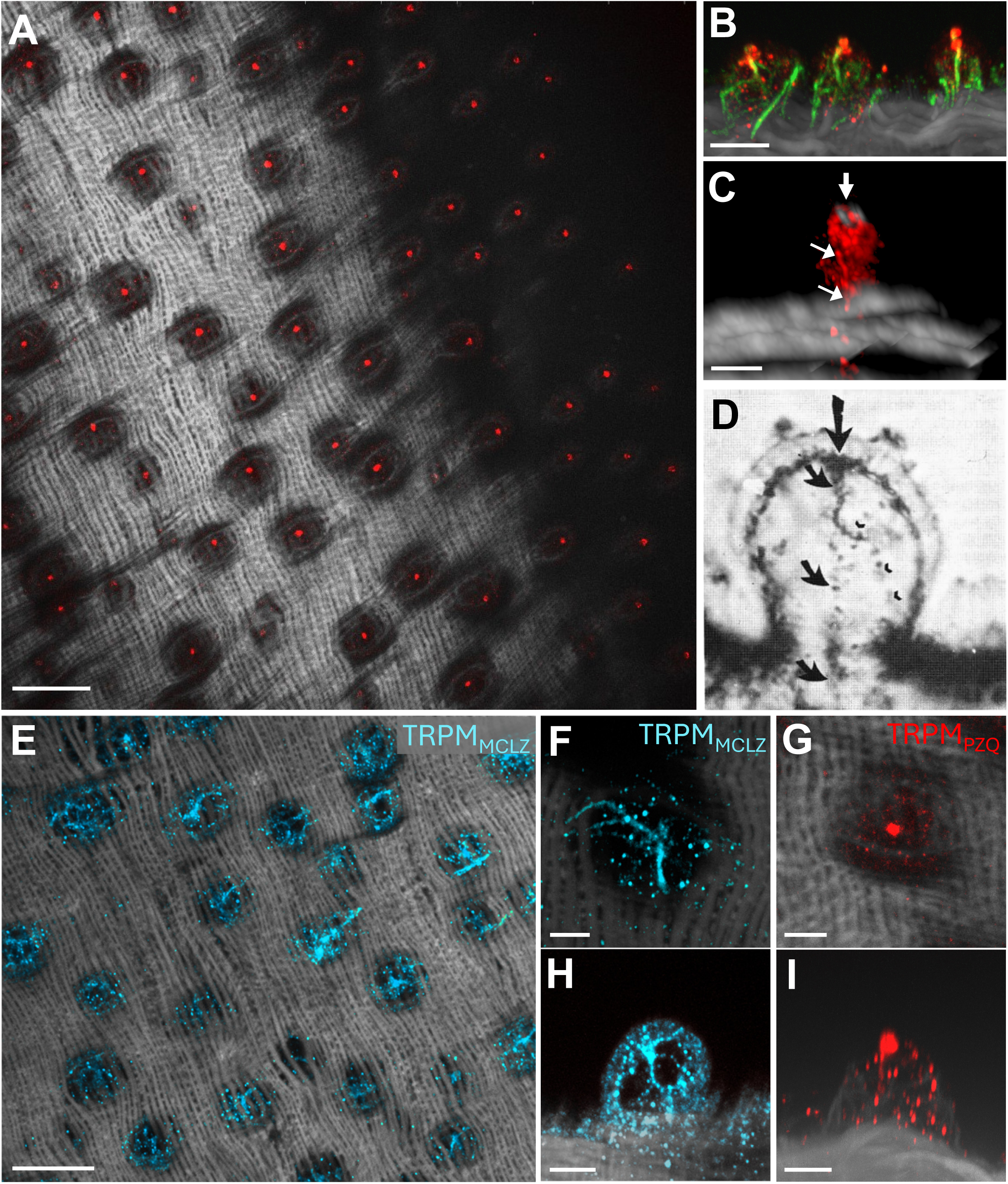
TRPM_PZQ_ localization in the tubercles of adult *S. mansoni* worms. (**A**) Surface view of TRPM_PZQ_ expression (red) within individual tubercles on the dorsal surface of an adult male worm. Grey, phalloidin staining. Scale bar: 20 μm. (**B**) Axial (‘xz’) projection of TRPM_PZQ_ localization (red) within tubercles, showing the intra-tubercular nerve network co-stained with β-tubulin (green). Scale bar: 5 μm. (**C**) 3D-STED image highlighting a corpuscle within a single tubercle. Scale bar: 2 μm. *Bold arrow* corpuscle, *smaller arrows* nerve process. (**D**) Image of a single tubercle stained for nerve tissue shown for comparison, reproduced without change from Kruger *et al.* 1986 (17). This work ©1986 through AOSIS Publishing is licensed under CC BY 4.0. To view a copy of this license, visit https://creativecommons.org/licenses/by/4.0/. *Large arrow,* terminal corpuscular process, *smaller arrows,* nerve supply to corpuscular process. (**E**) Surface view of TRPM_MCLZ_ expression (blue) in tubercles. Scale bar: 20 µm. (**F**) Magnified surface view of TRPM_MCLZ_ expression in processes within the tubercle. Scale bar: 5 µm. (**G**) Magnified surface view of punctate TRPM_PZQ_ expression (red) for comparison showing contrasting staining patterns. Scale bar: 5 µm. (**H**) Axial (‘xz’) projection of TRPM_MCLZ_ localization in a tubercle. Scale bar: 5 µm. (**I**) Axial (‘xz’) projection of TRPM_PZQ_ corpuscle for comparison. Scale bar: 5 µm.

The corpuscle is thought to be neuronal in nature, with a speculated chemosensory and/or mechanosensory role (24). In axial sections of individual tubercles, fluorescence from these TRPM_PZQ_-positive structures was contiguous with a β-tubulin-stained process that extended into the body of the worm (**Figure 3B**). The presence of multiple β-tubulin rich projections into the tubercle, not co-stained for TRPM_PZQ_, were suggestive of a more extensive intra-tubercular nerve network (Figure 3B). STED microscopy was utilized to obtain higher resolution images of the tubercle which captured a morphology resembling the interior of the corpuscular structure identified by Kruger & Hamilton-Attwell using electron microscopy (**Figure 3C&D,** Supplemental Video 1, (23, 24)).

TRPM_PZQ_ staining in the sensory corpuscle was positioned interior to a ring of actin marking the apex of the tubercle, as well as a process descending vertically into the parenchyma (Figure 3C&D). As a further control for the TRPM_PZQ_ staining pattern, we compared staining to that seen using an antibody targeting TRPM_MCLZ_ (12, 25).

TRPM_MCLZ_ was localized to discrete processes within the tubercle consistent with the broader intra-tubercular nerve network (**Figure 3E&F**) that was different from the sensory corpuscle staining seen with TRPM_PZQ_ (**Figure 3G**). This was also evident in axial (‘xz’) sections through an individual tubercle (**Figure 3H&I**).

TRPM_PZQ_-positive sensory papillae were particularly abundant in the area around the ends of the gynaecophoric canal (**Figure 4A**). At the upper end of the canal, TRPM_PZQ_ expression was present on both the left and the right edges (Figure 4A). Expression was primarily within hemispherical papillae and a fourth type of papillae, previously referred to as ‘macule’ papillae (14) that are found between the ventral sucker and the genital pore (Figure 4A, *inset*). As the male worm folds over onto itself to establish the gynaecophoric canal, a distinction can be made between the outer covering “flap”, which contains tubercles, and the inner non-tuberculated “flap.” On the outer flap, TRPM_PZQ_-positive hemispherical papillae were evident on both the interior and exterior edges along the length of the worm (**Figure 4B&C**). TRPM_PZQ_ staining was far less evident on the inner flap (Figure 4B&C, Supplemental Video 2). Within the gynaecophoric canal itself, TRPM_PZQ_ staining was absent, except for the interior distal end of the canal, where some channel expression was evident in papillae (Figure 4C).

**Figure 4.**
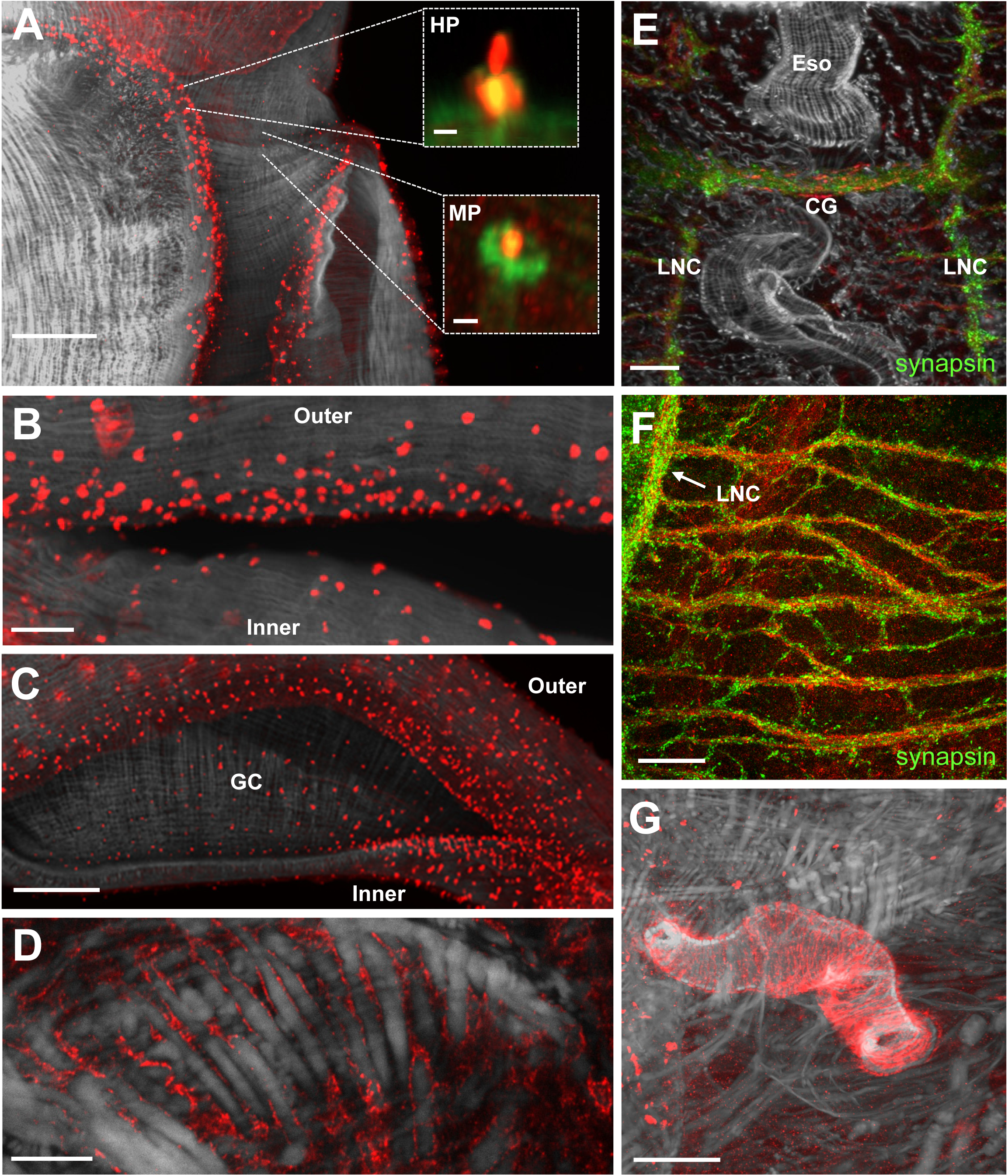
TRPM_PZQ_ staining within the body of adult *S. mansoni*. (**A**) Maximum intensity projection showing TRPM_PZQ_ localization on the “shoulders” of male *S. mansoni* surrounding the gynaecophoric canal. Scale bar: 50 μm. *Inset*: Magnified view of indicated regions to highlight TRPM_PZQ_ and β-tubulin within hemispherical (*top*, HP) and macule (*bottom*, MP) papillae. Scale bars: 2 μm. (**B**) Maximum intensity projection of TRPM_PZQ_ positive sensory endings on the outer (upper) flap of an adult male schistosome (head to left of sample). Scale bar: 20 μm. (**C**) Maximum intensity projection of TRPM_PZQ_ at the inferior terminus of the gynaecophoric canal (GC) showing the edges of both sides. Scale bar: 50 μm. (**D**) Confocal section interior to the tegument showing TRPM_PZQ_ within the sarcolemma of longitudinal muscle fibers. Scale bar: 10 μm. (**E**) Confocal section displaying TRPM_PZQ_ distribution in the cephalic ganglia (CG) and lateral nerve cord (LNC) projections co-stained with synapsin (green). Scale bar: 20 µm. (**F**) Confocal section from the worm mid region showing TRPM_PZQ_ localization to the peripheral nervous system, co-stained with synapsin (green). Scale bar: 20 µm. (**G**) TRPM_PZQ_ localization to the smooth muscle of the cirrus in males. Scale bar: 20 µm.

Throughout the entire body of the male adult worm, low levels of TRPM_PZQ_ staining was evident in the sarcolemma of longitudinal muscle fibers (**Figure 4D**). TRPM_PZQ_ was also found in the nervous system within the cephalic ganglia (**Figure 4E**), ventral nerve cords and the peripheral nervous system (**Figure 4F**). TRPM_PZQ_-positive peripheral processes predominantly extended ventrolaterally but also dorsolaterally at lower intensities. TRPM_PZQ_ was also strongly expressed in neural processes on both the left and right periphery of the oral sucker innervating the circular muscle fibers residing within the lower ‘lip’ (Supplementary Figure 6). Finally, TRPM_PZQ_ staining was observed in the reproductive tissues. In males, TRPM_PZQ_ localized to the circular muscle fibers that make up the cirrus, but no other region of the male reproductive system (**Figure 4G**).

TRPM_PZQ_ staining in adult female worms was of lower intensity but exhibited some similarity to that seen in adult males (**Figure 5A**), encompassing staining in nerve, muscle and sensory structures. TRPM_PZQ_ was found in digital form papillae throughout the head (**Figure 5B**) and hemispherical papillae on the surface of the body (**Figure 5C-D**). TRPM_PZQ_ expression in longitudinal muscle was maintained in females with expression throughout the body (Figure 5D). Central and peripheral nervous system staining for TRPM_PZQ_ was also evident. The channel was found in neuronal projections extending both to the periphery and in the commissural rings connecting the nerve cords (**Figure 5E**). TRPM_PZQ_ also localized to the female reproductive system. Staining was evident in the region of the Mehlis’ gland (**Figure 5F**) and the ootype (**Figure 5G**). TRPM_PZQ_ was found in the smooth muscle surrounding the distal end of the oviduct and the entry to the uterus. A TRPM_PZQ_-positive nerve network surrounding the exterior of the ootype was also evident (Figure 5G).

**Figure 5.**
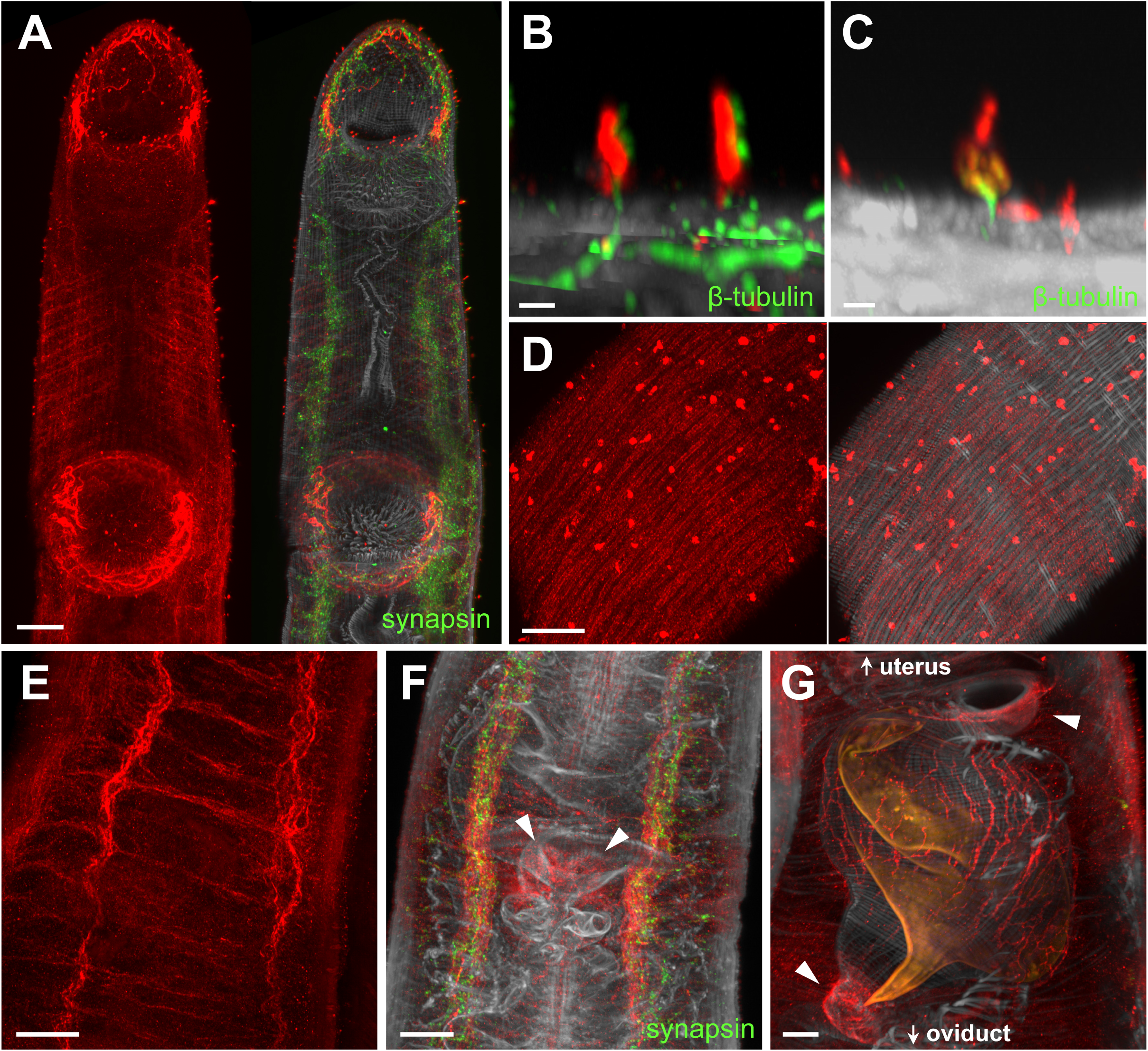
TRPM_PZQ_ localization in female *S. mansoni*. (**A**) Maximum intensity projection of staining within an adult (49 DPI) female *S. mansoni* worm using antibodies against phalloidin (grey), synapsin (green) and TRPM_PZQ_ (α-TRPM_PZQ_-N, red). Scale bar: 20 μm. (**B**) Digital form papilla found on the head of an adult female stained with TRPM_PZQ_ (red) and β-tubulin (green). Scale bar: 2 µm. (**C**) TRPM_PZQ_ and β-tubulin staining in a hemispherical papilla found on the body of a female worm. Scale bar: 2 µm. (**D**) TRPM_PZQ_ localization to the longitudinal muscle in adult females. *Left,* TRPM_PZQ_ alone; *right,* TRPM_PZQ_ and phalloidin staining. Scale bar: 20 µm. (**E**) Confocal section showing TRPM_PZQ_ localization to the nerve cords and commissures in female worms. Scale bar: 20 µm. (**F**) TRPM_PZQ_ staining in the Mehlis’ gland (white arrows). Scale bar: 20 µm. (**G**) TRPM_PZQ_ localization to the smooth muscle on either end of the ootype (white arrows) and within a nerve network surrounding the ootype. Autofluorescence of the developed egg (orange) aids identification of reproductive system components. Scale bar: 10 μm.

### Miracidia

TRPM_PZQ_ staining was evident across the entire miracidial surface in a pattern distinct from phalloidin and β-tubulin staining (**Figure 6A**). The TRPM_PZQ_ staining pattern decorated the ciliated epidermal plates but was absent from the epidermal ridges. In axial sections, TRPM_PZQ_ was localized externally to the muscle layer shown by phalloidin staining (**Figure 6B, *upper***). There was some immunofluorescence of TRPM_PZQ_ within the cilia, but the bulk of the staining localized internally to β-tubulin immunoreactivity in the cilia (**Figure 6B, lower**). In triple-stained samples, TRPM_PZQ_ exhibited clear separation between the muscle layer and the cilia (**Figure 6C**). This distribution suggests localization to the region surrounding the basal body of the cilia and the ciliary rootlet. Rootlets are enclosed in a membrane continuous with the epidermal plate plasma membrane and connect to the basal body of each cilium providing vital structural support (26). In some images colocalization of TRPM_PZQ_ and β-tubulin indicates that the channel is present in the basal body, yet in others a distinct separation would suggest that TRPM_PZQ_ is primarily found in the ciliary rootlet.

**Figure 6.**
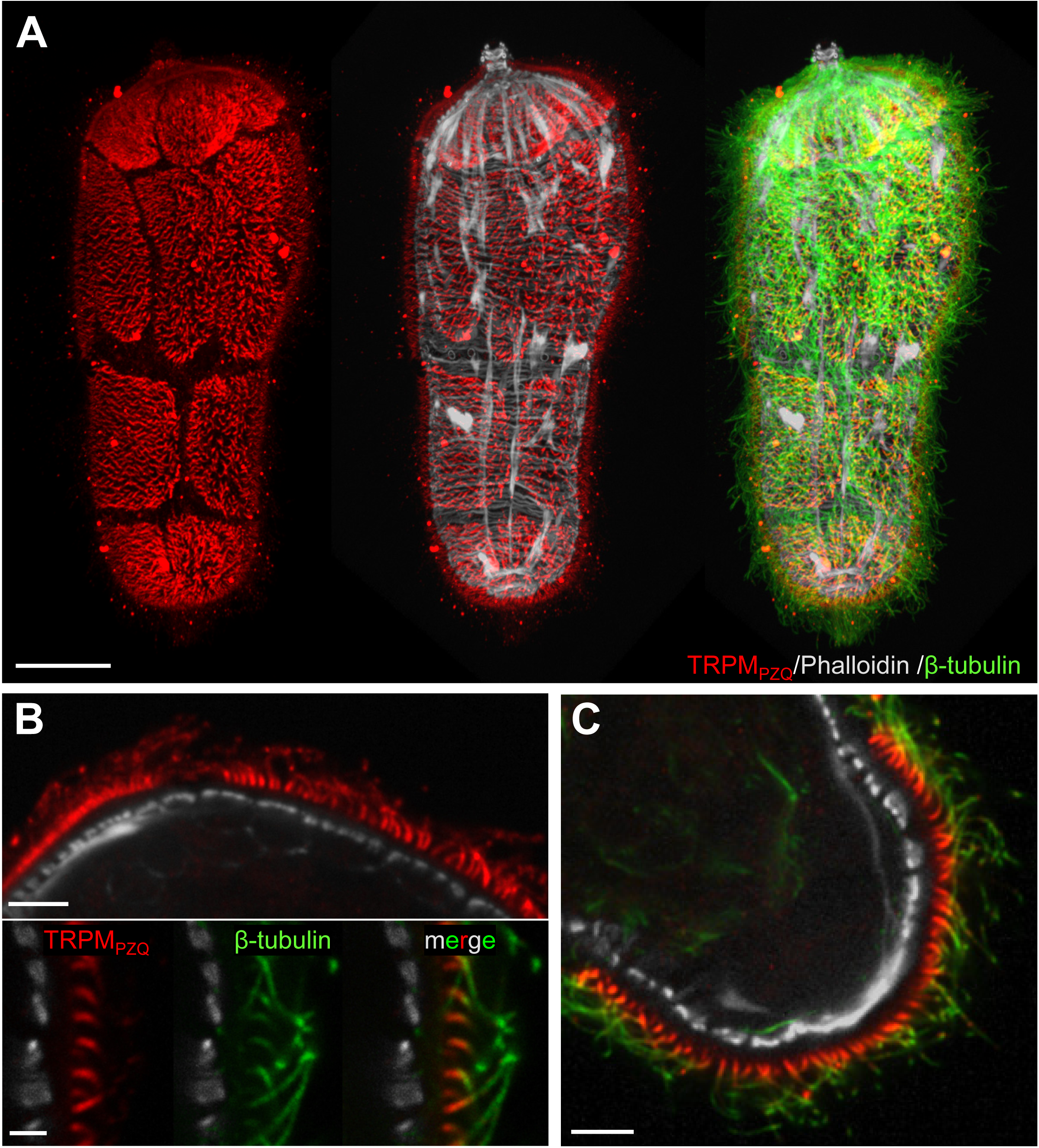
Expression of TRPM_PZQ_ in *S. mansoni* miracidia. (**A**) Maximum intensity projection of a miracidium stained with TRPM_PZQ_ (left), phalloidin (middle), and β-tubulin (right). Scale bar: 20 μm. (**B**) Localization of TRPM_PZQ_ to the putative ciliary rootlet and cilia. *Top,* TRPM_PZQ_ and phalloidin. Scale bar: 5 µm. *Bottom,* TRPM_PZQ_ and β-tubulin. Scale bar: 2 μm. (**C**) Immunostaining showing separation of the muscle layer (phalloidin, grey), rootlet (TRPM_PZQ_, red), and cilia (β-tubulin, green). Scale bar: 5 μm.

This is consistent with measurements of the TRPM_PZQ_ stained region that match recorded rootlet lengths (∼2 µm, (26)).

### Cercariae

TRPM_PZQ_ staining was evident in both the larvae body and the tail of cercariae (**Figure 7A**). Within the body, TRPM_PZQ_ was predominantly localized to nervous tissue (**Figure 7B**); the channel was found throughout the cephalic ganglia (**Figure 7C**) and within several neural processes (**Figure 7D**). Three main sets of neural processes were evident that were immunopositive for TRPM_PZQ_. These processes innervated (1) the pre acetabular/escape duct (PaD), (2) the periphery of the mouth (M), and (3) the ventral sucker (VS) (Figure 7D). TRPM_PZQ_ also localized to the short, rounded sensory papillae and the pairs of dorsal and ventral pits (27) that surround the gland openings at the apical tip of the body (**Figure 7E**). Within the tail, TRPM_PZQ_ was localized to a pair of gangliated nerves running parallel to the longitudinal nerve cords. These nerve cords spanned from the body-tail junction to the furca (**Figure 7F**).

**Figure 7.**
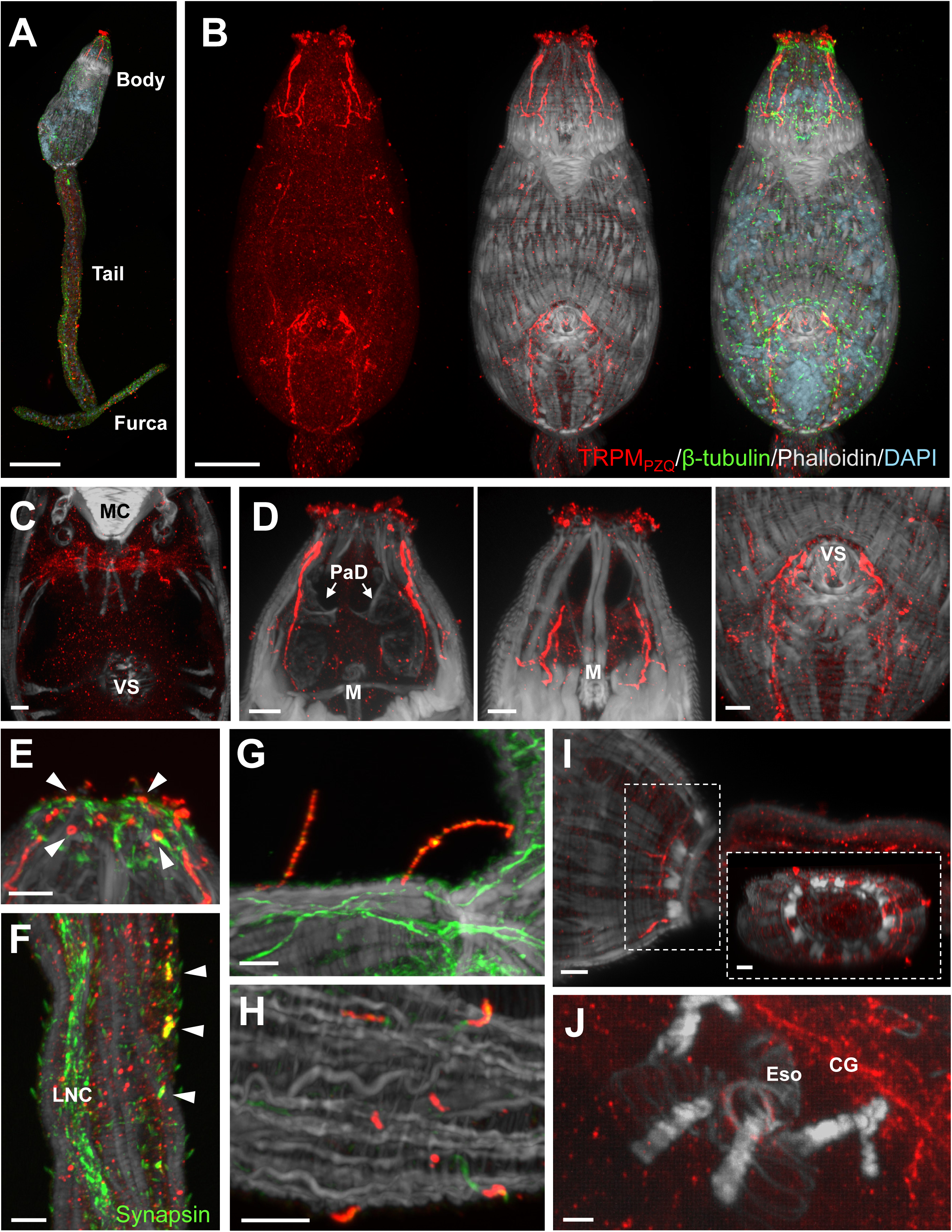
TRPM_PZQ_ localization to nervous and muscle tissue in cercariae. Unless otherwise noted, all cercariae samples were labeled with TRPM_PZQ_ (red), β-tubulin (green), phalloidin (grey) and DAPI (light blue). Abbreviations: muscle cone (‘MC’), ventral sucker (‘VS’), pre-acetabular ducts (‘PaD’), mouth (‘M’), longitudinal nerve cord (‘LNC’), esophagus (‘Eso’), cephalic ganglia (‘CG’). (**A**) Maximum intensity projection overview of TRPM_PZQ_ staining in a *S. mansoni* cercaria. Scale bar: 50 μm. (**B**) Maximum intensity projection of the body of the cercariae, showing TRPM_PZQ_ staining. Scale bar: 20 μm. (**C**) Confocal section showing TRPM_PZQ_ labeling of the cephalic ganglia. Scale bar: 5 μm. (**D**) TRPM_PZQ_ innervation of the: *left*, pre-acetabular ducts, *middle*, muscle cone, and *right* ventral sucker. Scale bars: 5 μm. (**E**) Maximum intensity projection of the sensory papillae and four pits (white arrows) surrounding the glands at the apex of the cercarial body. Scale bar: 5 μm. (**F**) Maximum intensity projection of the cercaria tail Showing TRPM_PZQ_ localization to dual gangliated nerve cords (arrowed) running parallel to the longitudinal nerve cord, labeled with synapsin. Scale bar: 5 μm. (**G**) long hair-like sensory projections along the tail, stained with TRPM_PZQ_ and β-tubulin. Scale bar: 5 μm. (**H**) Shorter uniciliate TRPM_PZQ_-positive sensory papillae on the furca labeled with TRPM_PZQ_ and β-tubulin. Scale bar: 5 μm. (**I**) Maximum intensity projection of TRPM_PZQ_ labeling of the muscle surrounding the body-tail junction. *Inset,* cross-sectional view of the body-tail junction. Scale bars: 5 μm. (**J**) Confocal section showing TRPM_PZQ_ localization to the smooth muscle at the junction of the esophagus and the bifurcated gut. Scale bar: 2 μm.

TRPM_PZQ_ was also found within different types of sensory structures along the tail. These included long hair-like cilia found along the tail (**Figure 7G**) and shorter uniciliate papillae found on the furca (**Figure 7H**). Finally, TRPM_PZQ_ staining was also evident in two distinct muscle groups in cercariae: the first being musculature at the junction of the body and the tail surrounding muscle fibers that make up the base of the body (**Figure 7I**); second, at the terminal end of the esophagus, immediately preceding the bifurcation of the primordial gut (**Figure 7J**).

### Schistosomula

The localization of TRPM_PZQ_ in 24-hour schistosomula was similar to the distribution described within cercarial body (**Figure 8A**). The TRPM_PZQ_ innervation of the muscle cone and ventral sucker was maintained, as was expression in the cephalic ganglia (**Figure 8B**). TRPM_PZQ_ staining was evident in the uniciliate sensory papillae distributed throughout the body of the schistosomulum (**Figure 8C**). TRPM_PZQ_ staining was again apparent in the papillae surrounding the gland openings but was more restricted in schistosomula, localizing to newly developed uniciliate papillae rather than the smaller rounded papillae bodies and pits seen in cercariae. These papillae varied in number between samples (**Figure 8D**), which may reflect different stages of transformation and/or development. TRPM_PZQ_ was expressed at the basal end of the schistosomula, surrounding the musculature of the former body-tail junction (**Figure 8E**). Finally, TRPM_PZQ_ staining was consistently observed in the gut, surrounding the smooth muscle of the posterior sphincter of the esophagus (**Figure 8F**).

**Figure 8.**
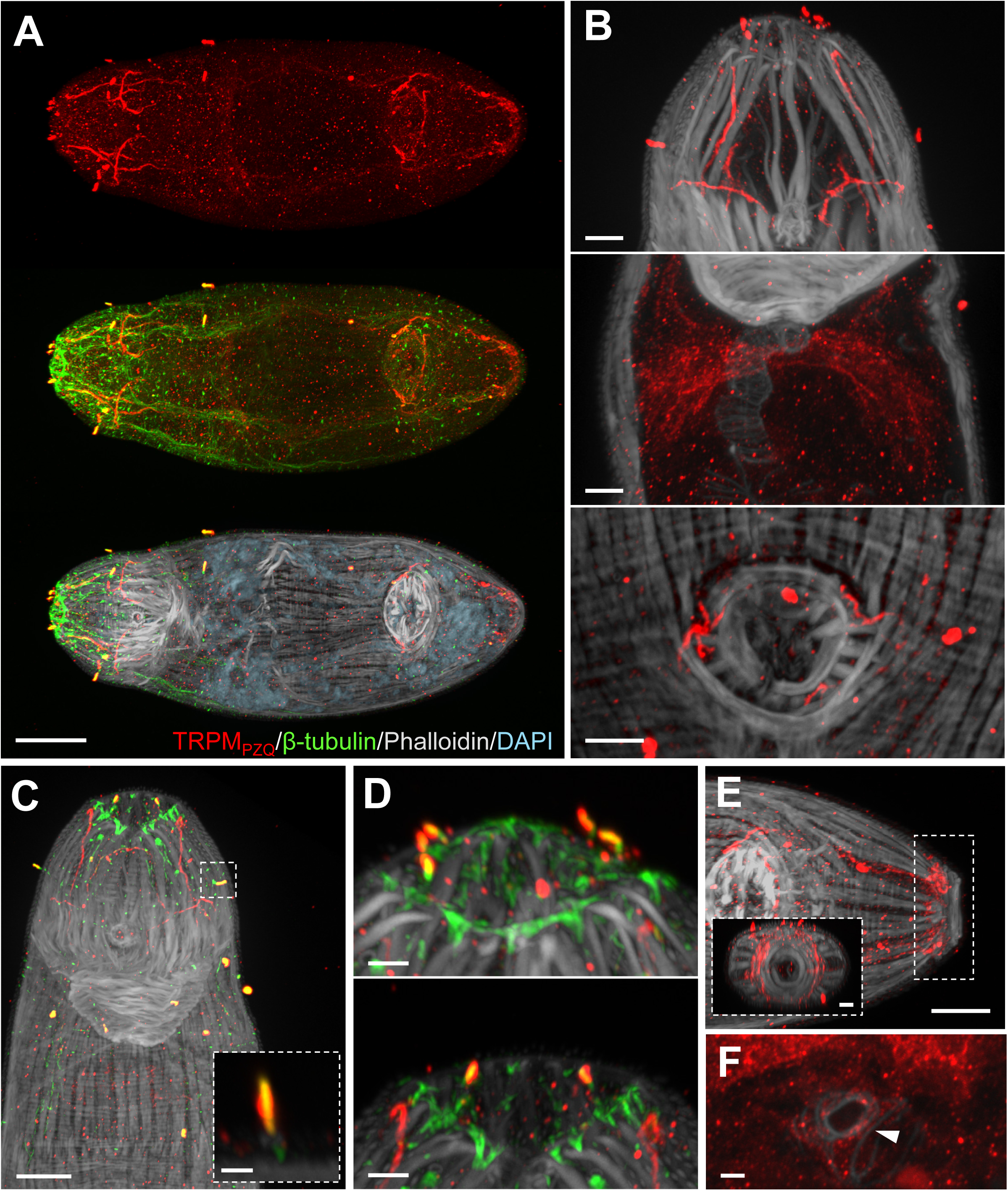
TRPM_PZQ_ localization in 24h schistosomula. (**A**) Maximum intensity projection overview of TRPM_PZQ_ localization. All samples were labeled with TRPM_PZQ_ (red), β-tubulin (green), phalloidin (grey) and DAPI (light blue). Scale bar: 20 μm. (**B**) Confocal section showing labeling of neural projections anterior to the muscle cone (*top*), cephalic ganglia (*middle*), and ventral sucker (*bottom*). Scale bars: 5 μm. (**C**) Maximum intensity projection showing TRPM_PZQ_ labeling of sensory papillae. *Inset* magnified view of boxed region, displaying morphology. Scale bar: 10 μm, inset scale bar: 2 μm. (**D**) Maximum intensity projection of apical tip of schistosomula. TRPM_PZQ_ is localized to the sensory papillae, which varied in number (top versus bottom) between samples. Scale bars: 2 μm. (**E**) Localization of TRPM_PZQ_ to the musculature surrounding the basal end of the schistosomula, as previously seen in cercariae. *Inset* side view. Scale bars: *main* 10 μm, *inset* 5 μm. (**F**) Confocal section of the gut, showing TRPM_PZQ_ localization to smooth muscle. Scale bar: 2 μm.

### TRPM_PZQ_ immunostaining in samples that display altered sensitivity to PZQ

We were interested to compare TRPM_PZQ_ expression and localization in samples known to exhibit differential sensitivity toward PZQ – first, a PZQ-susceptible (PZQ-ES) and PZQ-resistant (PZQ-ER) strain of *S. mansoni*, and second, examination of TRPM_PZQ_ staining in juvenile and adult worms.

First, we examined a PZQ-susceptible (PZQ-ES) and PZQ-resistant (PZQ-ER) strain of *S. mansoni* that have been generated from a laboratory strain of *S. mansoni* (LE).

These lines differ in their recovery to PZQ exposure by ∼377-fold (14, 28), Western blot analyses of adult male worm lysates revealed expression of TRPM_PZQ_ (∼250kDa) in both lines but with lower expression of TRPM_PZQ_ in the PZQ-ER line (**Figure 9A**), Densitometry of multiple blots (n=3) comparing these lines showed that expression of

**Figure 9.**
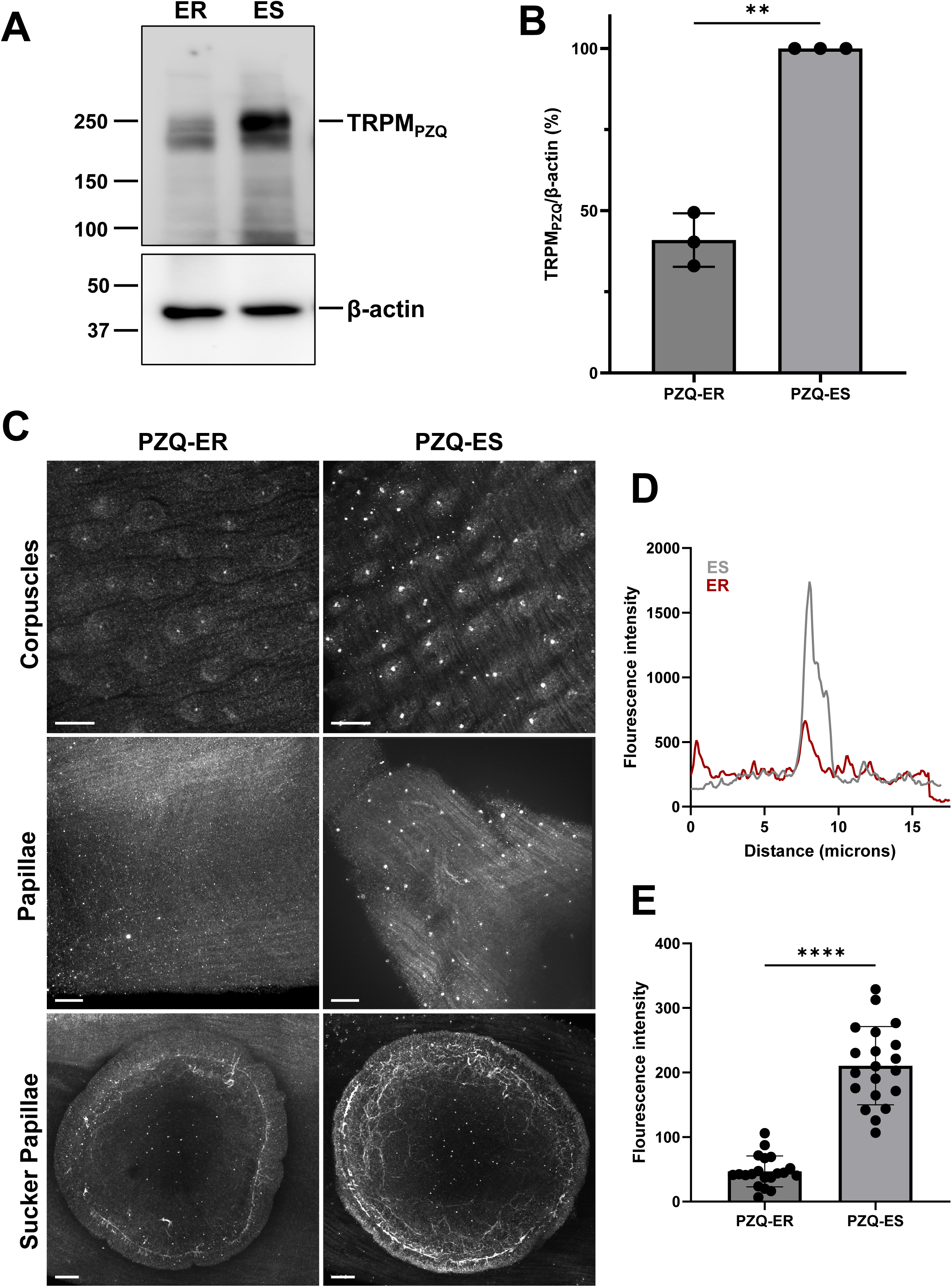
Comparison of TRPM_PZQ_ expression in PZQ resistant and susceptible *S. mansoni*. (**A**) Western blot of PZQ-ER (resistant) vs PZQ-ES (sensitive) strains in adult male worm lysate. (**B**) Densitometric measurements from independent blots in PZQ-ER and PZQ-ES males (n = 3). (**C**) Staining for TRPM_PZQ_ in PZQ-resistant (left) and PZQ-susceptible (right) worms. (**D**) Corpuscle fluorescence profile in PZQ-ER and PZQ-ES worms. Profile is from a single ∼20 µm line spanning a single tubercle. (**E**) Quantification of tubercle corpuscle intensity difference between PZQ-ER and PZQ-ES strains. Data are from 20 different tubercles spread across two male worms. Significance was determined for (**B**) and (**E**) using a Welch’s t-test, ** p < 0.01, **** p < 0.0001.

TRPM_PZQ_ in the PZQ-ER line was 50±4.8% compared with immunoreactivity in the PZQ-ES line (**Figure 9B**). Immunolabeling of TRPM_PZQ_ in both the lines was also performed. The intensity of immunostaining was lower in PZQ-ER compared with PZQ-ES adult worms, also indicative of decreased TRPM_PZQ_ expression. For example, the intensity of sensory corpuscle staining in tubercles was significantly lower in images captured from PZQ-ER compared with PZQ-ES worms (**Figure 9C**). This was quantified by measuring the peak intensity of fluorescence staining (**Figure 9D&E**). Similarly, sensory papillae found in the region between the two suckers showed reduced staining intensity in PZQ-ER worms compared with PZQ-ES worms (Figure 9C). TRPM_PZQ_ expression in the circular neuronal ring around the perimeter of the ventral sucker was also lower. Therefore, the overall trend was one of decreased intensity of TRPM_PZQ_ staining in the PZQ-ER strain.

Next, we examined TRPM_PZQ_ staining in juvenile worms which are known to display higher tolerance to PZQ. Here, we utilized the standardized reference stages of schistosome development as defined originally by Basch (29) to categorize the developmental stages. A staging key, modified from this publication (29), is shown in **Figure 10A**. Stages (x-axis) are based on mean worm length (y-axis) which broadly correlate with the infection durations annotated on the schematic (Figure 10A). In the context of development within the mouse model, we investigated ‘juvenile’ stages from the early liver stage (stages 5-7, ∼15 days post infection, 15 DPI) to the mesenteric stage (stage 18+, ∼28 DPI) during which worms pair and lay eggs (30–34). During this time, schistosomes undergo major development and morphological changes as illustrated in **Figure 10B**. In all images presented, a staging index is shown in the top right corner of each panel.

**Figure 10.**
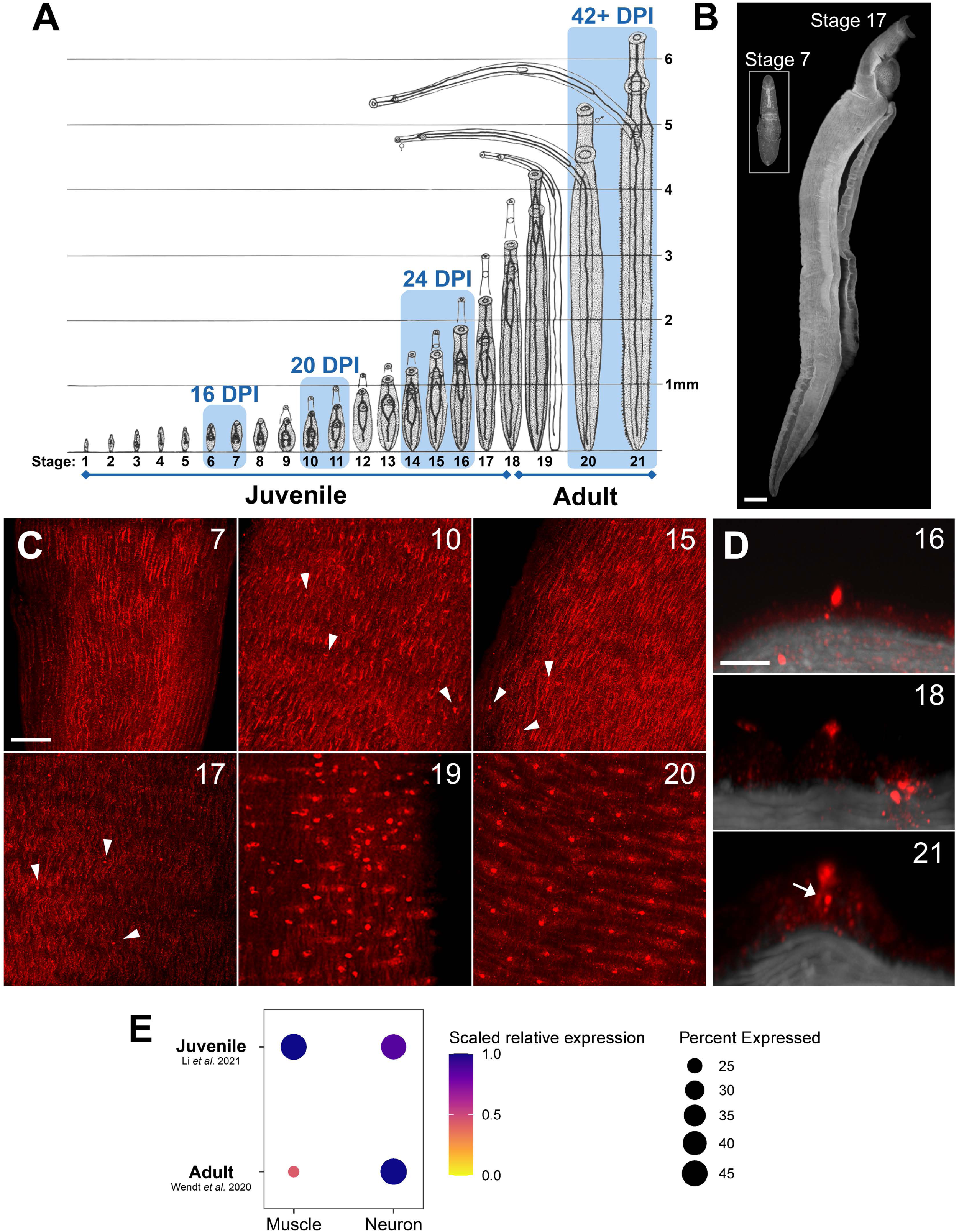
TRPM_PZQ_ expression during development of juvenile worms. (**A**) Staging of *S. mansoni* development in the murine host illustrating developmental stages (‘x’-axis) and mean worm (‘y’-axis, length in millimeters). Blue boxes correspond to timepoints post infection. Modified with permission from (22). (**B**) Image of stage 7 juvenile and stage 17 juvenile labeled with phalloidin to highlight the growth and development that occurs between these stages. Scale bar: 100 μm. (**C**) Overview of TRPM_PZQ_ localization changes between stage 7 and stage 20 male worms. TRPM_PZQ_ localized to longitudinal muscle fibers in the most immature worms, with development of sensory papillae only evident from stage 10 onwards (white arrows). Tubercle development began around stage 18. Scale bar: 20 μm. (**D**) Development of TRPM_PZQ_ staining in tubercles. The ‘xz’ projections show the increased size and volume of a tubercle with development and the extension of a TRPM_PZQ_ neural projection (arrow) connecting the bulb and the body. Each panel was captured from a different sample. Scale bar: 5 μm. Developmental stages are indicated in the top right corner of each panel. (**E**) Dot plot showing distribution of TRPM_PZQ_ mRNA in juvenile and adult *S. mansoni* worms collated from single cell RNA sequencing datasets from juvenile (17) and adult worms (18). Analysis of these datasets is described in the methods.

Unlike staining patterns identified in adult male *S. mansoni*, TRPM_PZQ_ was strongly expressed in peripheral muscle tissue across juvenile developmental stages. Longitudinal muscle staining was evident throughout the worm body at the earliest investigated stage (**Figure 10C**). This muscle staining was maintained throughout early juvenile stages but decreased in intensity as the tegument thickened and tubercles developed. At later maturation stages, TRPM_PZQ_-positive sensory papillae began to appear across the surface of both males and females concomitant with a decrease in expression of TRPM_PZQ_ in the subtegumental muscle (Figure 10C). From stage 18 onward, TRPM_PZQ_ localized to both the digital-form papillae interspersed between tubercles and the developing sensory corpuscle in each tubercle. TRPM_PZQ_ was expressed in these structures as they emerged from the surface with staining increasing in size and complexity as the tubercles matured (**Figure 10D**). By stage 21, when the tubercle was fully formed, TRPM_PZQ_ expression resided within the sensory corpuscle as well as the neuronal projection that extended down towards the muscle layer (Figure 10D). These data demonstrate a shift in TRPM_PZQ_ immunostaining from the peripheral muscle into neurons throughout juvenile worm development. In contrast, neuronal staining of TRPM_PZQ_ deeper within the worm was maintained throughout worm maturation from early into late-stage juveniles with expression evident in the ventral nerve cord and associated projections (Supplementary Figure 7).

### Comparison with single cell RNAseq datasets

We considered whether the observed transition of TRPM_PZQ_ from peripheral muscle to sensory structures during worm maturation might be a technical artifact from reduced antibody accessibility into worm tissue during development of the tegument. We considered this unlikely as staining of nervous tissue deep within the parenchyma of both juvenile and adult worms was consistently observed. However, independent insight can be gleaned from analysis of single cell RNA sequencing (scRNAseq) data. Therefore the distribution of TRPM_PZQ_ mRNA was queried in two published scRNAseq datasets for adult (18, 35) and juvenile (3.5 weeks, (17)) *S. mansoni* as described in the ‘Methods’. In both datasets, TRPM_PZQ_ mRNA was distributed across multiple muscle and neuronal cell clusters, which when collapsed into generic ‘muscle’ and ‘neuron’ categories displayed a similar pattern of bias toward localization in muscle cells in juveniles, but neurons in adult worms (**Figure 10E**). Therefore, the redistribution of TRPM_PZQ_ from a ‘muscle’ to ‘neuron’ localization during juvenile to adult worm development that was observed by immunofluorescence was also seen in the scRNAseq data.

In summary, this study details the expression pattern of TRPM_PZQ_ across multiple life cycle stages of *S. mansoni*, evidencing expression in muscle, nerve and various discrete sensory structures across the worm surface. In two different cases where schistosomes display decreased PZQ sensitivity, changes in TRPM_PZQ_ expression and/or localization were evident.

## Discussion

This study resolves TRPM_PZQ_ expression in nerve, muscle and sensory structures across various *S. mansoni* life cycle stages. While expression of TRPM_PZQ_ in various excitable cell types is unsurprising given known aspects of PZQ action - for example, PZQ-evoked spastic paralysis (4) and prior recording of TRPM_PZQ_-like currents in neurons (36) – a detailed spatial mapping of TRPM_PZQ_ localization furthers our understanding of this important drug target. These data help (i) interpretation of phenotypes evoked by PZQ exposure, (ii) deciphering the role(s) of TRPM_PZQ_ throughout the parasite life cycle, and (iii) demonstrating how changes in TRPM_PZQ_ localization and/or expression impact PZQ action. These topics are elaborated upon in the following three sections.

### TRPM_PZQ_ localization aligns with PZQ-evoked phenotypes

Resolving the expression pattern of TRPM_PZQ_ helps understanding the basis of PZQ-evoked phenotypes (11). For example, when adult worms cultured *in vitro* are exposed to PZQ, they typically detach from their culture vessels and unpair. TRPM_PZQ_ expression in the neuronal ring that circumscribes the ventral sucker (Figure 2B), and in the sensory structures innervating the muscle flaps that envelop the gynaecophoric canal (Figure 4), are consistent with these phenotypic observations. Expression of TRPM_PZQ_ in tubercles (Figure 3) is consistent with early damage to these structures evoked by PZQ (37, 38). In miracidia, low concentrations of PZQ evoke irregular swimming patterns with swimming inhibited at higher PZQ exposures (3–5). This impact on swimming behavior is also consistent with the localization of TRPM_PZQ_ to ciliary rootlets (Figure 6), where PZQ-evoked TRPM_PZQ_ activity will disrupt coordination of ciliary beating. Treatment of cercariae with PZQ also causes cessation of swimming, owing to rapid movements and contraction of the body and tail that precede separation of the tail from the body at the highest exposures (39–41). Activation of TRPM_PZQ_ in sensory papillae and the gangliated nerve cords (16) found within the tail and furca likely underpin the rapid tail movements evoked by PZQ (Figure 7). TRPM_PZQ_ in the musculature surrounding the body-tail junction in cercariae is the likely effector for body/tail detachment after PZQ treatment, or during host penetration (Figure 7). Lastly, PZQ treatment induces emptying of the penetration gland in a *Trichobilharzia* species (42), and TRPM_PZQ_-positive innervation is clearly evident in the pre-acetabular duct (Figure 7) where it is well placed to cause exocytotic Ca^2+^ fluxes that mediate secretion. Therefore, from these multiple examples, the localization of TRPM_PZQ_ to particular cells and structures aligns with phenotypic effects evoked by PZQ.

### TRPM_PZQ_ expressed in various sensory structures across worm surface

Confocal microscopy revealed expression of TRPM_PZQ_ in a diverse range of sensory structures across the worm surface. These sensory structures have previously been detailed from studies using electron microscopy (21–24). That TRP channels are found within sensory structures is entirely consistent with their role as environment sensors, as TRP channels have been shown to detect natural ligands and/or other environmental cues – such as light, temperature, pH, oxidative stress and mechanical forces (43). The presence of TRPM_PZQ_ in these sensory structures infers a likely chemosensory and/or mechanosensory role for TRPM_PZQ_, which could vary between the different structures and different life cycle stages. Successful completion of the parasite life cycle necessities transitioning between distinct environments, and TRPM_PZQ_ and other schistosome TRP channels may help guide these transitions between different environments. Such a role would underscore vulnerability to chemotherapeutic attack and we note that the two schistosome TRPM channels deorphanized to date are both activated by anthelmintic chemotypes (PZQ and BZQ for TRPM_PZQ_ (8, 44), meclonazepam and M4339 for TRPM_MCLZ_ (12, 45)).

While the native role of TRPM_PZQ_ has yet to be defined, prior work has shown that TRPM_PZQ_ can be activated by membrane stretch (46). Resolution of TRPM_PZQ_ localization in these sensory structures could therefore imply a mechanosensitive role. For example, TRPM_PZQ_ expression in and around the ciliary rootlet in miracidia is consistent with the role of the rootlet in transducing mechanical stimuli or sensing mechanical stress (47–49). Rootlets are found in both motile and non-motile cilia and couple to other structures within the cilia in order to convey forces to mechanosensitive ion channels (50, 51). The localization of TRPM_PZQ_ in the sensory corpuscle of tubercles could also contribute to mechanosensation of adult worms, possibly as a sensor for blood pressure (46), directionality of blood flow, or contact with the blood vessel lining, or other worms. Preliminary functional imaging of these structures suggests a basal neuronal tone within tubercles, which may reflect natural movements and distension of the worm surface. We visualized the presence of other structures (nerve fibers, cytoskeletal structures) and ion channels (TRPM_MCLZ_) within tubercles, suggesting a multisensory role for these structures (Figure 3). The immunoprecipitation dataset for TRPM_PZQ_ also identified spectrin (found in tubercles (52)) and paramyosin (also found on the worm surface (53, 54)), potentially indicative of mechanosensitive coupling.

Similarly, TRPM_PZQ_-positive sensory structures at the termini of the gynaecophoric canal may fulfill a mechanosensory role in sensing the presence of a female worm and regulating pairing/unpairing. We certainly do not exclude chemosensory roles for TRPM_PZQ_ in any of these sensory structures – as, consistent with the polymodal nature of TRP channel activation, it is possible that TRPM_PZQ_ senses multiple types of stimuli. A good analogy is found by considering the sensory biology of different types of ‘taste’ buds found on our tongue, which impart chemosensitivity to different flavors as well as mechanosensitive roles in feeling food texture, with often both sensations being transduced in the same type of structure (55). One can imagine that surface sensory structures such as the TRPM_PZQ_-positive papillae on the surface of both suckers may allow the worm to ‘feel’ (‘mechanosensitivity’) and ‘taste’ (‘chemosensitivity’) their local environment.

The sensory role of the surface TRPM_PZQ_-positive structures is also intriguing for other schistosome life cycle stages. In cercariae, the localization of TRPM_PZQ_ to papillae and pits surrounding the gland duct and innervation of the pre-acetabular gland is likely relevant for host recognition and penetration (Figure 5D&E). Similarly, the role of TRPM_PZQ_-positive papillae and cilia along the tail (Figure 5G&H) merits further study in the context of chemotaxis and host-seeking behavior. Further effort to understand the roles of these structures would be worthwhile given their likely functional importance to parasite physiology.

### Changes in TRPM_PZQ_ expression and localization correlate with changes in schistosome sensitivity to PZQ

We studied TRPM_PZQ_ in two different scenarios where changes in worm sensitivity to PZQ are apparent. First, we examined PZQ-ER and PZQ-ES adult worms (Figure 9). Second, we compared TRPM_PZQ_ localization and expression between juvenile and adult worms (Figure 10).

The *S. mansoni* enriched populations, PZQ-ER and PZQ-ES, derived by Le Clec’h exhibited a 377-fold difference in viability after PZQ exposure *in vitro* (14). This difference in PZQ sensitivity between these strains correlated with a ∼2.25-fold decrease in levels of TRPM_PZQ_ mRNA in PZQ-ER compared with PZQ-ES worms (14). Western blotting of these same lines revealed a lower expression of full length TRPM_PZQ_ in PZQ-ER compared with PZQ-ES (Figure 9A). The measurements of mRNA (56% decrease (14)) or protein (59% decrease, Figure 9A) in the PZQ-ER population are in close agreement and demonstrate decreased expression of TRPM_PZQ_ protein in PZQ-ER correlates with higher PZQ tolerance. The lower the amount of TRPM_PZQ_ expressed, the smaller the PZQ-evoked current will be, providing a higher capacity for recovery from PZQ exposure. If a ∼60% decrease in TRPM_PZQ_ expression correlates with a ∼400-fold change in tolerance toward PZQ exposure (14), then monitoring TRPM_PZQ_ expression levels in natural strains of schistosomes must be a key component of surveillance efforts aimed at monitoring PZQ efficacy in natural worm populations (56).

TRPM_PZQ_ is expressed in juvenile worms, as shown by Western blotting for the full-length protein (Figure 1G&H). This result is consistent with observations that both adult and juvenile worms rapidly contract in response to PZQ (5). Why then do juvenile worms show higher tolerability to PZQ exposure than adults (5, 57–59)?

The explanation for this resilience may be multifactorial. For example, it has been proposed that PZQ is more actively exported from juvenile than adult worms (60). Juvenile worms are growing rapidly and presumably have higher regenerative potential than adults (17, 61), which would provide greater capacity for repairing PZQ-evoked damage. However, the observations of (i) a slightly lower expression of TRPM_PZQ_ protein in juveniles compared with adult males (Figure 1G&H) and (ii) the observed transition of TRPM_PZQ_ from a ‘muscle’ to ‘neuron’ localization during juvenile to adult schistosome development (Figure 10) are also likely relevant. In terms of expression, the lower levels of TRPM_PZQ_ in juveniles will result in a lower PZQ-evoked current and less of a toxic insult. In terms of localization, it is likely that TRPM_PZQ_-evoked Ca^2+^ signals are better tolerated in muscle. This is because muscle cells, unlike neurons, display high capacity for Ca^2+^ buffering. In vertebrates, the voluminous sarcoplasmic reticulum (SR) in muscle contains specialized buffers such as calsequestrin that possess low affinity but high capacity for calcium binding. Consequently, muscle SR can sequester large quantities of Ca²_⁺_ (up to ∼ 20 mM total (62)). This high storage capacity mitigates any risk of cellular Ca²_⁺_ overload, while still supporting the large cyclic Ca²_⁺_ fluxes necessary for rapid, repetitive muscle contraction. In contrast, neurons typically exhibit lower capacity for Ca^2+^ storage, utilizing buffers such as calreticulin which possess moderate affinity and lower Ca^2+^ buffering capacity. Neurons have a known vulnerability to Ca^2+^ overload (63, 64), and the deleterious effects of excitotoxic Ca^2+^ levels will be propagated to neighboring cells through dysregulated release of neurotransmitters. As PZQ activation of TRPM_PZQ_ mediates a large, sustained depolarizing current (65), the Ca^2+^ buffering capacity of different cells is likely a critical factor in keeping free Ca^2+^ below a level that is toxic. While schistosome muscle is morphologically distinct from muscle types in vertebrates (66), it is a reasonable inference that activation of TRPM_PZQ_ in neurons (adult worms) will be more detrimental to the worm than activation of TRPM_PZQ_ in muscle (juvenile worms). This explanation is consistent with prior observations that PZQ treatment of juvenile worms does not cause extensive damage to subtegumental tissues (57), despite the presence of TRPM_PZQ_ in muscle. Rather, the extent of PZQ-evoked tegumental damage increases during worm development as expression of TRPM_PZQ_ transitions to surface sensory structures. We also note the lesser efficacy of PZQ on adult female worms compared with male worms may also relate to the considerably lower expression of TRPM_PZQ_ in females (Figure 1G&H), as well as the greater distribution of TRPM_PZQ_ in muscle in female compared with adult male worms (Figure 5D).

In conclusion, immunostaining of TRPM_PZQ_ revealed expression in various sensory structures, as well as nerve and muscle tissue. Changes in TRPM_PZQ_ expression (PZQ-ER vs PZQ-ES) and/or tissue localization (juvenile vs adult schistosomes) correlated with known scenarios of PZQ tolerance.

## Acknowledgements

Schistosome infected mice were provided by the NIAID Schistosomiasis Resource Center at the Biomedical Research Institute (Rockville, MD) through NIH-NIAID Contract HHSN272201000005I for distribution via BEI resources. All imaging was completed at the Oxford Instruments Center for Advanced Microscopy - Electron Microscopy Core at the Medical College of Wisconsin (RRID:SCR_026315). STED imaging was performed using an Abberior Facility Line STED microscope, which is supported by the NIH S10 Award S10OD034247-01A1 (to Dr. Xuelin Lou) and managed by Oxford Instruments Center for Advanced Microscopy - Electron Microscopy Core. This work was supported by the NIH (R01-AI145871 to JSM, F31-AI183573 to CMR).

Supplementary Information (Supplementary Figures x 7, Videos x 2)

