## Supplementary Material for "Expression of TRPM_PZQ_ in excitable cells and sensory structures throughout the *Schistosoma mansoni* lifecycle"

Rohr *et al*.

Supplementary Information (Figure x 7, Video x 2)

**Supplementary Figures**

**Supplemental Figure 1. Identification of Sm.TRPM$_{PZQ}$.** Sequence of Sm.TRPM$_{PZQ}$ highlighting peptides identified in endogenous immunoprecipitation assays from male worms as shown in Figure 1D. Peptides highlighted in yellow were found in an excised 250kDa band from immunoprecipitations performed with the $NH_2$-terminal antibody (α-TRPM$_{PZQ}$-N), but not the control (IgG) immunoprecipitation performed in parallel.

**Supplemental Figure 2. Western blot of exon-deleted TRPM$_{PZQ}$ constructs.** *Top,* Constructs of *Sm*.TRPM$_{PZQ}$ differing by omission of single exons (noted above each lane) were used to validate epitope specificity of the $NH_2$-terminal TRPM$_{PZQ}$ antibody (α-TRPM$_{PZQ}$-N) via Western blot. Deletion of exon 4 resulted in a loss of immunoreactivity. Bottom, β-actin is shown as the loading control.

**Supplemental Figure 3. Secondary only antibody controls.** (**A-F**) Images of different lifecycle stages that were stained with the secondary antibody alone. These control samples were processed in parallel with other experimental samples but the primary antibodies (TRPM$_{PZQ}$, β-tubulin, or synapsin) were omitted. All samples were incubated in secondary antibodies (either goat-anti-rabbit Alexa Fluor 568, or goat-anti mouse Alexa Fluor 488), fluorophore conjugated phalloidin, and DAPI. Different life cycle stages are labelled as indicated.

**Supplemental Figure 4. A TRPM$_{PZQ}$-positive ring of neurons runs alongside circular muscle surrounding the ventral sucker.** Confocal section of the circular muscle ring on the rim of the oral sucker stained with phalloidin (grey, top) or TRPM$_{PZQ}$ (red, middle) with a merged composite (bottom). Scale bar: 10 µm.

**Supplemental Figure 5. Comparison of tubercle staining with both $NH_2$-terminal and COOH-terminal antibodies.** The TRPM$_{PZQ}$ antibodies generated in this study (α-TRPM$_{PZQ}$-N, left; α-TRPM$_{PZQ}$-C, right) both recognize the corpuscle structure at the center of each tubercle. Scale bar: 20 µm.

**Supplemental Figure 6. TRPM$_{PZQ}$ localization in the oral sucker of an adult male schistosome.** Maximum intensity projection of the oral sucker of *S. mansoni* (*left*) stained with phalloidin (grey), β-tubulin (green), and TRPM$_{PZQ}$ (red), scale bar: 10 µm. *Right,* innervation of longitudinal muscle fibers in the oral sucker (taken from the region in the main panel indicated by the dashed boxes) and overlayed with TRPM$_{PZQ}$ and β-tubulin staining. Scale bar: 10 µm.

**Supplemental Figure 7. Development of TRPM$_{PZQ}$ staining in juvenile worms.** (**A**) Side view of a maximum intensity projection showing TRPM$_{PZQ}$ (red) staining in a stage 13 juvenile worm co-stained with phalloidin (grey). TRPM$_{PZQ}$ is present in both muscle and neural tissue. Scale bar: 50 µm. (**B**) Changes in TRPM$_{PZQ}$ staining in muscle layer between the suckers across stage 6, stage 10, and stage 17 juveniles. Scale bars: 20 µm. White arrows indicate the development of sensory papillae. (**C**) A deeper confocal section within the same stage 13 juvenile worm as shown in panel A demonstrating $TRPM_{PZQ}$ staining in the central nervous system. Scale bar: 50 μm. (**D**) $TRPM_{PZQ}$ staining and the development of the central and peripheral nervous system during juvenile maturation highlight the development of lateral projections (white arrows) and connectives between the ventral and dorsal nerve cords (white star). Scale bars: 20 μm.

**SF1**

MTEQIPFQRTSTLKHRISRTGSVGQDNNNNITGSINISSGINTGIQQQSSIRRASVMQKYMGEI
EFTGLNQTARFCKLNSETPDSILRDILKRKWGLKPPTLIITVFGTDFEKKRKLKMIFKKGLWKA
AESGCWIVTGGFHLGVMKLTGEAVRDYTDAYGGNRMMAFGVASWDCVTKNEILEAALHEGTAVY
QSEEDEEEEAEDSEPILVDPLSQGGIKSVRSDIEERALDPNHNFFVLVDDGTTDQLKGKEAELR
ARFERCISLWNCASSPQEQTTTVTSQQSGPISSTTHLQQPQNGISTSSNVNNNVNLGGLQRQGS
TMGGQASTTSGPQTSATSIGKTSAITGGSSSNSALSKGVSEPVVSNQPVSTASVTRGTKDKLTT
ESSLKPAKSGKSVGSEEEILVPMCGLVVGGDRFTLRQVYCSMIRNRCPIVVTKGTSGAADVLAF
GLDAANKMASEEVVDDKDQVPLEARLESIIEEFLGDLHPDYTNYTDEVNMLCEIINEYTNLVSV
FDMEEDSDLDGYVISSLLASAGSTVASDQINLEQLEITLTLNRADIAREKIFLENKKWKKGQLN
DYMYQALMSDRHDFVKLFLEQGFSLEDFLTIYMLEKLYTDQLKNLNSKVAIFNKMWEYNRSHRT
SKVALRDVGKVIKALVGDFYHPLYLSKEFQAKLMPEKRLEGADICQRLLGSNKLRSSQDYSNMS
PSSSSSSTSSSSHHNGAICPRTLGLKDNVRFIDYNEDYDLMNLNSLHLTSPPPPLPPPPPTSTT
TVIEEIGFRNPAFLECDKYGTRAIIRRGATSVEPAEKNDGDNDDDDNDYDVDDREDHDDDNEIY
GSNMNGFTMNSRLPRSSTTMTTVSGEPLISSNVLRYNDKGNTMGTSTVSHLGSSRIRDGTIQQQ
YHLNDEGNSSRPHNTDDMSSQLKKSKYLSATRNNDNGNRNGYSSRNKHGRRPSSQTNEHDMLIP
GQPLIYNATITNINRSGHSSRGPYTSLGPGHFRITGRAPLAYDATSMGTTTPIQSPKPIINDTL
QFSFDNEHVQRNSVLISSIPNEKSNKPVLFEDETINKPSRSCLMSCIQRITSFCSRLFRCNTDT
KKVNKGLSLSQESTGIKGPKEFATLRAMAALAAATAATAVADPTKTTQGDLIIPSNFFEPDEEQ
TKSIQLDRPARELLIWSILVGKLRMAELFWTMEKEPIAAALLASILLTSLGNKTDDFTDKEDYR
SFAKNFQERAEGVLNECYREDEHRTQLIINHELIYYGRSSVIKLAAEGQSIKFMAHPCCQDFLT
NTWSGNLSTKNSVFRYIMGVVCGLTLPFLIPKVILSKPKTNPTTEGEESSPTSEQQGSTQSNHT
MKYVLTAEDAGKKGVRKTLQARTVEYISQIRDFYMAPVVRFVYNTISYITFLILFSYLLLVDFR
INITVVEYIVIAWVITLFIEEIKQIAWAVLSGISFRTYISDGWNKLDCAGLALYIIGFILRLIV
LLRLGQENNFNIKYERYYIVTDPILDPSRICLAISLFTFYIRLMYTFSFHIALGPKLIMIGKMV
TNDLIPFMIILTVIMVGYAVAAQSIAYPNGLFTKENMTLNSEIKHMTFADIIFSMYTTAYFQMF
GDFSLDALQGEDRTCQNHMCPTKTSRWLVPIMLGFYVLLTNILMFNLLIAMFSKTYEEIESAST
YYWNYQRYQMINDYVHRSPLAPPIIIVWHFYEAYKAIGNQCASLRNAETAKYNPFCVRFTDVKK
EREMVKWEHMKAMDYLREPSTKATGKRGVAESRAVVFRGGGGVGPGQGPVMDLKSEMSSVTEGI
GMELEKRFKEIDNQFQRFNDVDSRLNDVTQLLTNLSEVISNVTETQQRIIRQINELPTCQCNEF
TDEVVPVQKSTTHQTDSGTAETRKRTKLEVAIQSALEAAKDVLRPPTPPPPPPPPPPPQGKESP
VLLAAVVPGSEDDSSGSEEGGDPSADPVLVPQVPDIKTGRVIERTLGDHRLWRMAPFNFEKYPG
MRMNVPPERMAWTVEYPDYFAYDACEEILLFPSEESHDGVNFTHGNIVFTFQHSENLRTINFNQ
YDAKAKLRRQSLLGRYRLDSTTGAPLNPMGRTGLLGKGLLPRWGPNHSFVLCITRWTRDTRTGA
QIIRSNRGVLQYLALERNKRLCMPWYLTDHTSKCDFDECVLKIISSLITRRGHAMLPEKRVDRL
LKRIEKAEVTQIFKGYLDDQLNADSAWMETVVINLHESESKGAQLPDDILKLLNEPNTEEQSKW
IEVSHSSNLRTSHNYILKNVAELRRAFY

SF2

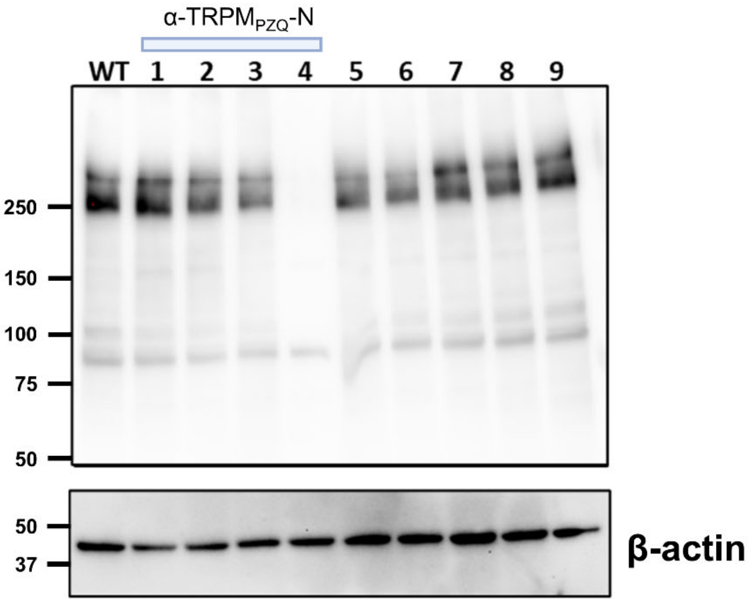

SF3

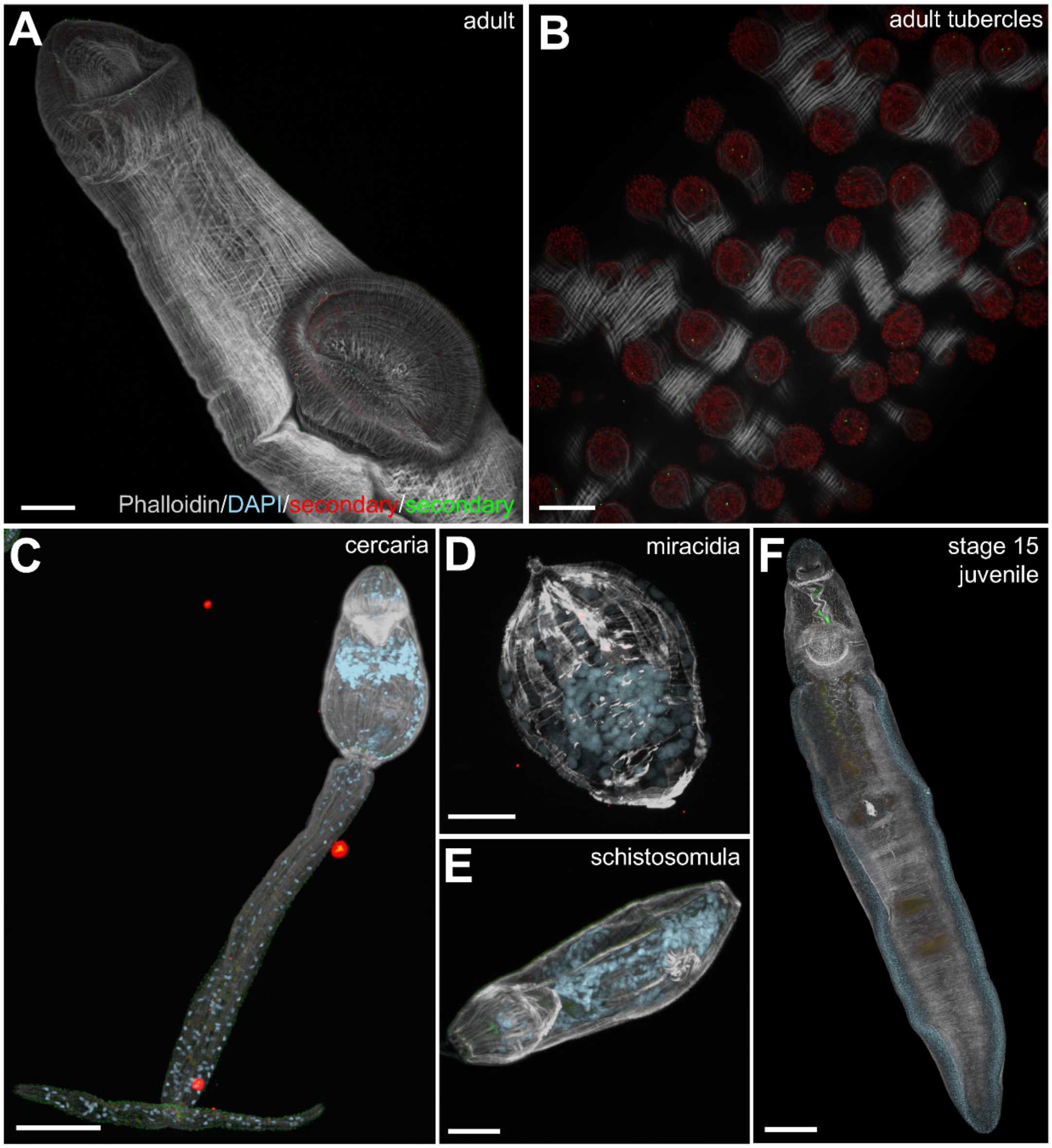
A
adult
Phalloidin/DAPI/secondary/secondary
B
adult tubercles
C
cercaria
D
miracidia
E
schistosomula
F
stage 15
juvenile

**SF4**

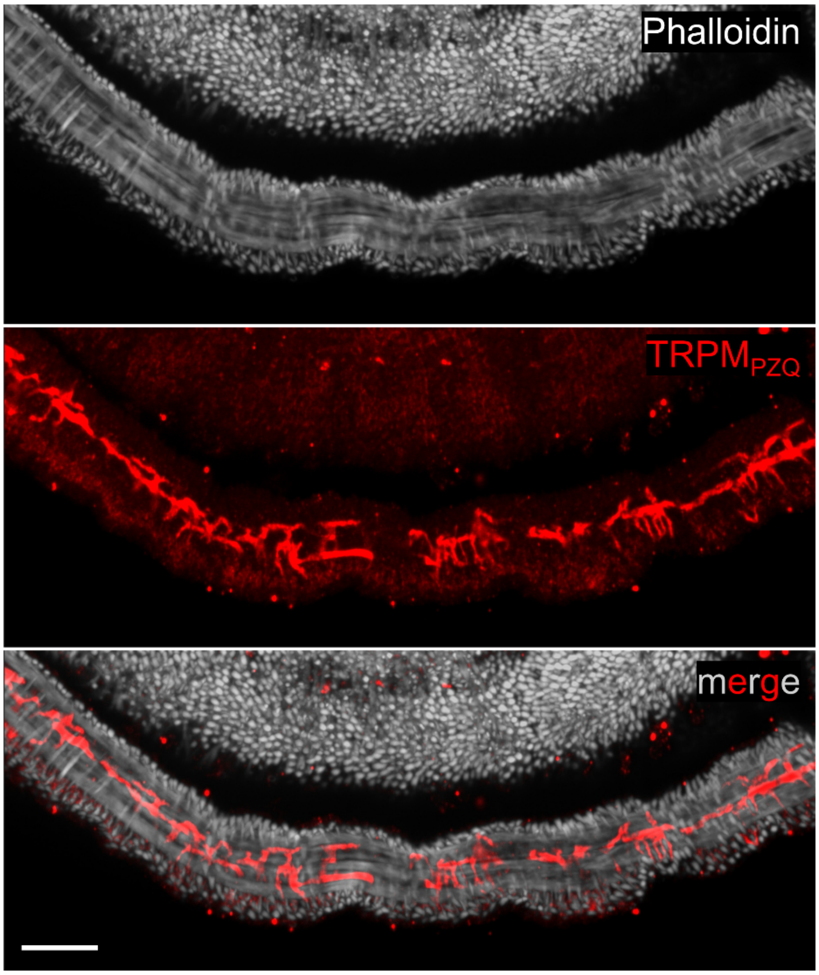

SF5

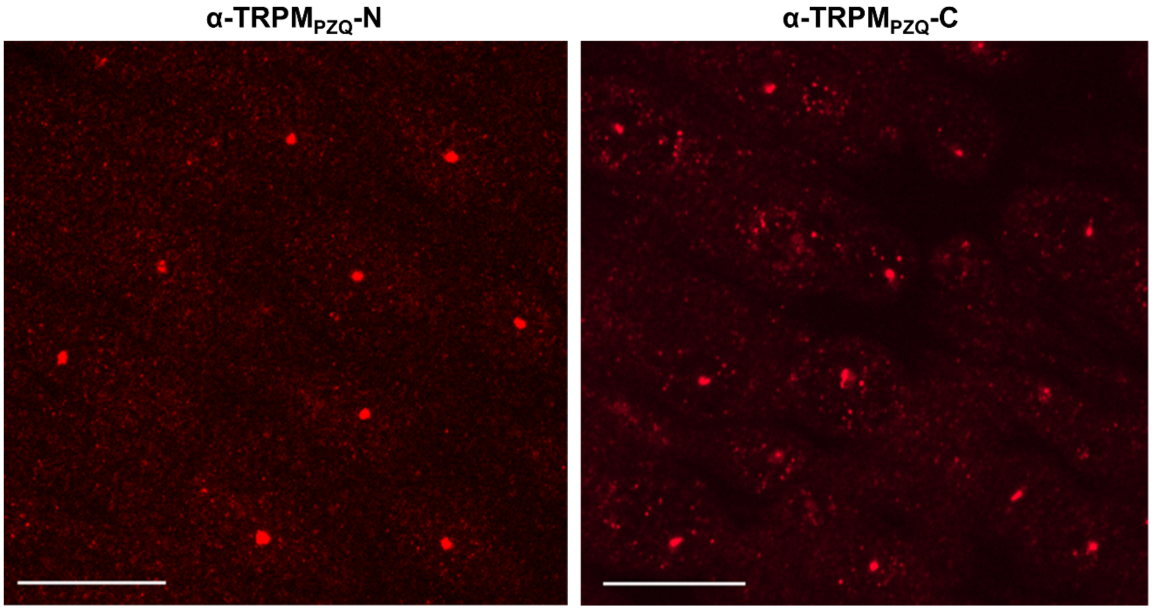

**SF6**

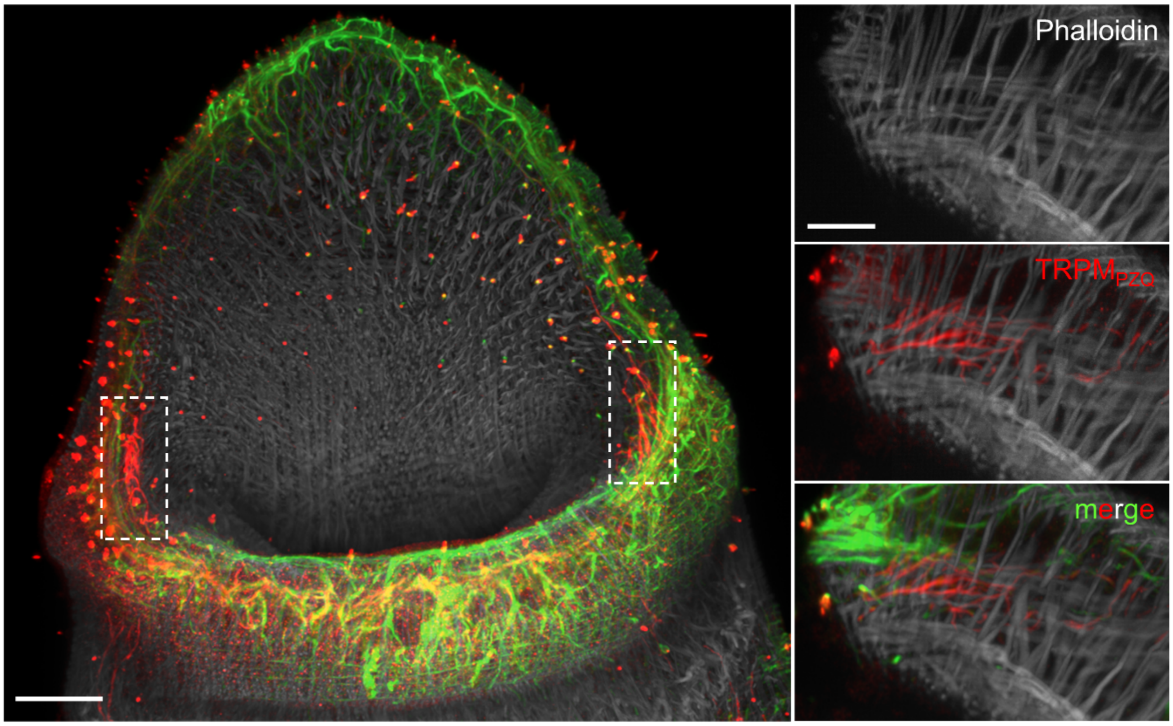

**SF7**

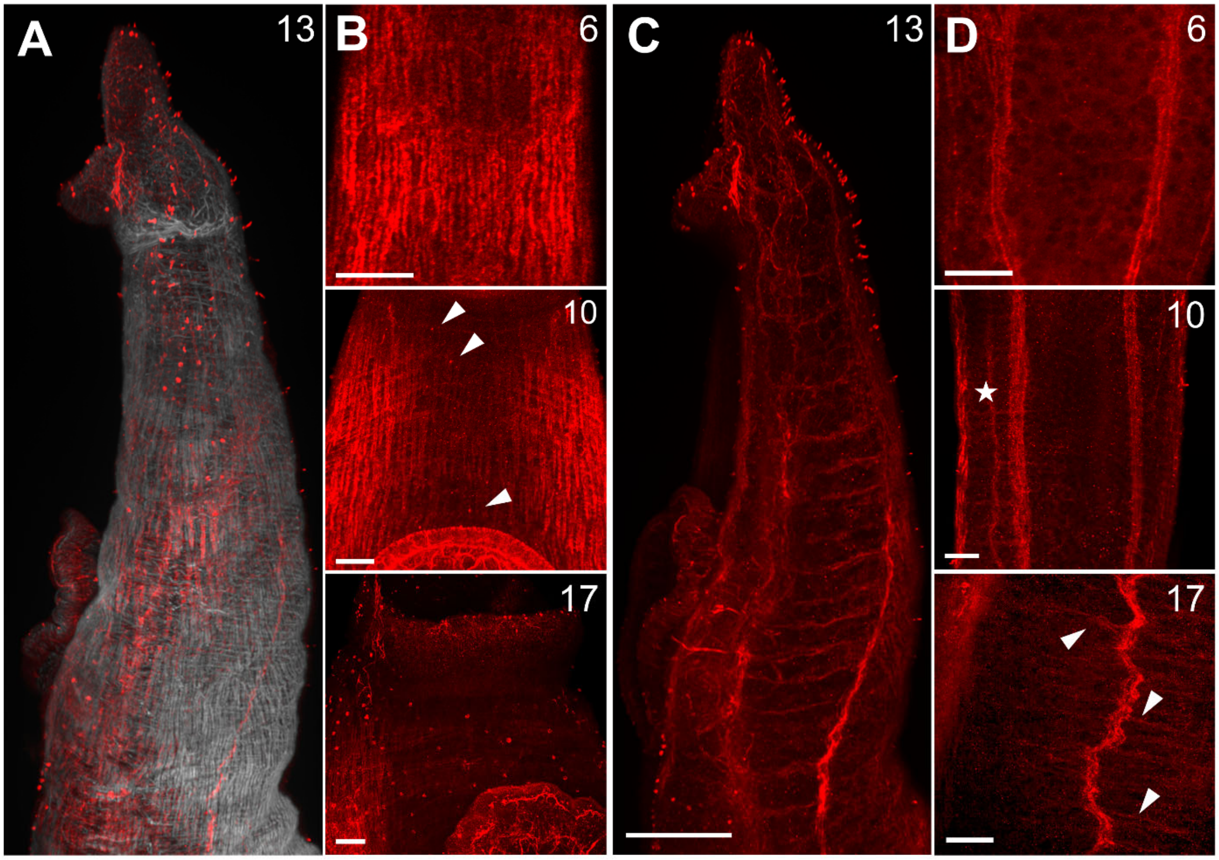

### Supplementary Videos

**Supplementary Video 1. $TRPM_{PZQ}$ in the sensory corpuscle.** 3D rendering (360° rotation) of a tubercle labeled with phalloidin (grey), β-tubulin (green), and $TRPM_{PZQ}$ (red).

**Supplementary Video 2. Distribution of sensory papillae on the edges of the gynaecophoric canal.** 3D rendering showing the asymmetric distribution of $TRPM_{PZQ}$-positive sensory papillae on the outer and inner flaps of an adult male worm. Staining is evident on both the exterior and interior of the outer flap.
